# Quantifying the influence of genetic context on duplicated mammalian genes

**DOI:** 10.1101/2025.04.03.647042

**Authors:** Alexander S. Moffett, Andrea Falcón-Cortés, Michele Di Pierro

## Abstract

Gene duplication is a fundamental part of evolutionary innovation. While single-gene duplications frequently exhibit asymmetric evolutionary rates between paralogs, the extent to which this applies to multi-gene duplications remains unclear. In this study, we investigate the role of genetic context in shaping evolutionary divergence within multi-gene duplications, leveraging microsynteny to differentiate source and target copies. Using a dataset of 193 mammalian genome assemblies and a bird outgroup, we systematically analyze patterns of sequence divergence between duplicated genes and reference orthologs. We find that target copies, those relocated to new genomic environments, exhibit elevated evolutionary rates compared to source copies in the ancestral location. This asymmetry is influenced by the distance between copies and the size of the target copy. We also demonstrate that the polarization of rate asymmetry in paralogs, the “choice” of the slowly evolving copy, is biased towards collective, block-wise polarization in multi-gene duplications. Our findings highlight the importance of genetic context in modulating post-duplication divergence, where differences in cis-regulatory elements and co-expressed gene clusters between source and target copies may be responsible. This study presents a large-scale test of asymmetric evolution in multi-gene duplications, offering new insight into how genome architecture shapes functional diversification of paralogs.

**Significance statement:** After a gene is duplicated, reduced selective constraints can lead the two copies to rapidly diverge, with one copy often evolving faster and occasionally gaining a new function. We quantify the influence of genetic context in choosing which copy of a duplicated gene has an elevated substitution rate. In a representative dataset of 193 mammalian genomes, we found strong evidence that gene copies pasted into new genomic locations tend to evolve faster than the corresponding copies in ancestral locations, suggesting an important role for the regulatory environment. The asymmetry in evolutionary rates of duplicated genes persists even for very large multigenic duplications, up to the scale of megabases, indicating that regulatory interactions frequently reach farther than previously thought.

---

Gene duplications play a crucial role in evolution, providing frequent opportunities for functional innovation. If a gene duplication fixes in a population, both copies can face relaxed purifying selection, leading to loss of function in one of the copies [1, 2]. However, several mechanisms have been proposed to explain long-term maintenance of both copies, including neofunctionalization, subfunctionalization, escape from adaptive conflict, and selection for increased gene dosage [3]. Most of the possible fates of a duplication are expected to result in asymmetric evolution of the two copies, as has been widely observed [4, 5, 6, 7, 8, 9, 10, 11, 12, 13, 14, 15, 16, 17, 18, 19]. While it is not immediately clear which of the two copies will experience an accelerated rate of evolution, previous work suggests that the copy in the ancestral genomic location tends to evolve at similar rate before and after the duplication, while the copy inserted into a new location often experiences an accelerated evolutionary rate. Adopting the terminology of Dewey [20], we refer to the copy in the original location as the source copy and the the relocated copy as the target copy. If the target copy is more likely to have a higher substitution rate than the source copy, we say that asymmetrically evolving paralogs are polarized. Polarization of paralogs has been shown in small collections (varying between one and six) of model mammalian species [8, 9, 10, 13, 15] and nematodes [12]. Depending on the specific duplication event, this increase of evolutionary rate in the source copy can be consistent with positive selection [9, 13] or relaxed negative selection.

Substitution rate asymmetry between paralogs is often location-dependent, highlighting differences in the regulatory environment of the source and target copies, where cis-regulatory elements and coexpressed genes can be drastically different depending on the location of a gene [19]. Polarization has also been found to correlate with local recombination rate in yeast [6], suggesting that the relevant genetic context cannot be reduced to only cis-regulatory elements. Regardless of the specific nature of genetic context, both the distance between copies and the size of duplicated regions are factors that might influence polarization of paralogs. Both of these attributes of duplications reflect the conservation of genetic context between paralogs, where closeby source and target copies are likely to directly share cis-regulatory elements while large duplications may carry cis-regulatory elements and co-expressed genes to the source location. Paralogs resulting from tandem duplications do not seem to evolve asymmetrically [8, 10]. In *C. elegans*, the genomic distance between target and source copies on the same chromosome has little effect on rate asymmetry, while relocation of the target copy to a different chromosome is associated with accelerated evolution [12]. In the same study, the length of the duplicated region was inversely related to rate asymmetry, so that smaller duplications tend to evolve more asymmetrically. Additionally, asymmetric evolution in multi-gene duplications is not well-characterized, despite the importance of segmental duplications [21, 22, 23, 24, 25, 26, 27, 28, 29, 30] and whole-genome duplications [31, 32, 33]. Paralogs resulting from whole-genome duplications, called “onhologs” or “homeologs”, have been found to evolve asymmetrically [7, 34], while a study on multi-gene duplications in *A. thaliana* did not find significant asymmetric evolution [35].

Despite considerable progress in characterizing the polarization of gene duplications, more work is needed in order to uncover quantitative relationships among inter-paralog distances, sizes of duplicated regions, and rate asymmetry and polarization. Here, we address this gap in the literature, analyzing gene duplication in a broad sample of mammals. We focus on multi-gene duplications, which allows us to reliably identify source and target copies and quantitatively analyze the effects of genetic context on polarization. We test the location-dependence of paralog polarization using multi-gene duplications found through microsynteny analysis. Using these duplications and estimates of their duplication time, we find strong evidence for polarization, where source copies tend to evolve slower than target copies. We observe decreasing polarization with the size of duplicated regions and a weak relationship between polarization and the genomic distance between the two copies on the same chromosome, while paralogs on different chromosomes exhibit the strongest polarization. Finally, we observe block-wise polarization for simultaneously duplicated genes. Altogether, these results highlight the importance of genetic context in the evolutionary fate of gene duplications.

We first identify microsynteny blocks, regions of conserved gene order, by analyzing the structure of gene homology across pairs of genomes. Microsynteny forms a basis for comparing genomes of distantly related species and accurately inferring orthology [36, 37, 38, 39, 40, 41, 42, 43, 44]. We use microsynteny to identify both source and target copies (Fig. 1a), where the largest microsynteny block that a gene belongs to (primary microsynteny) is likely the source copy. The correspondence between primary microsynteny and source copies follows from the likelihood that the flanking genes around the duplicated region have conserved order in species diverging before the duplication event, while the target copy is will not have this property.

**Figure 1:**
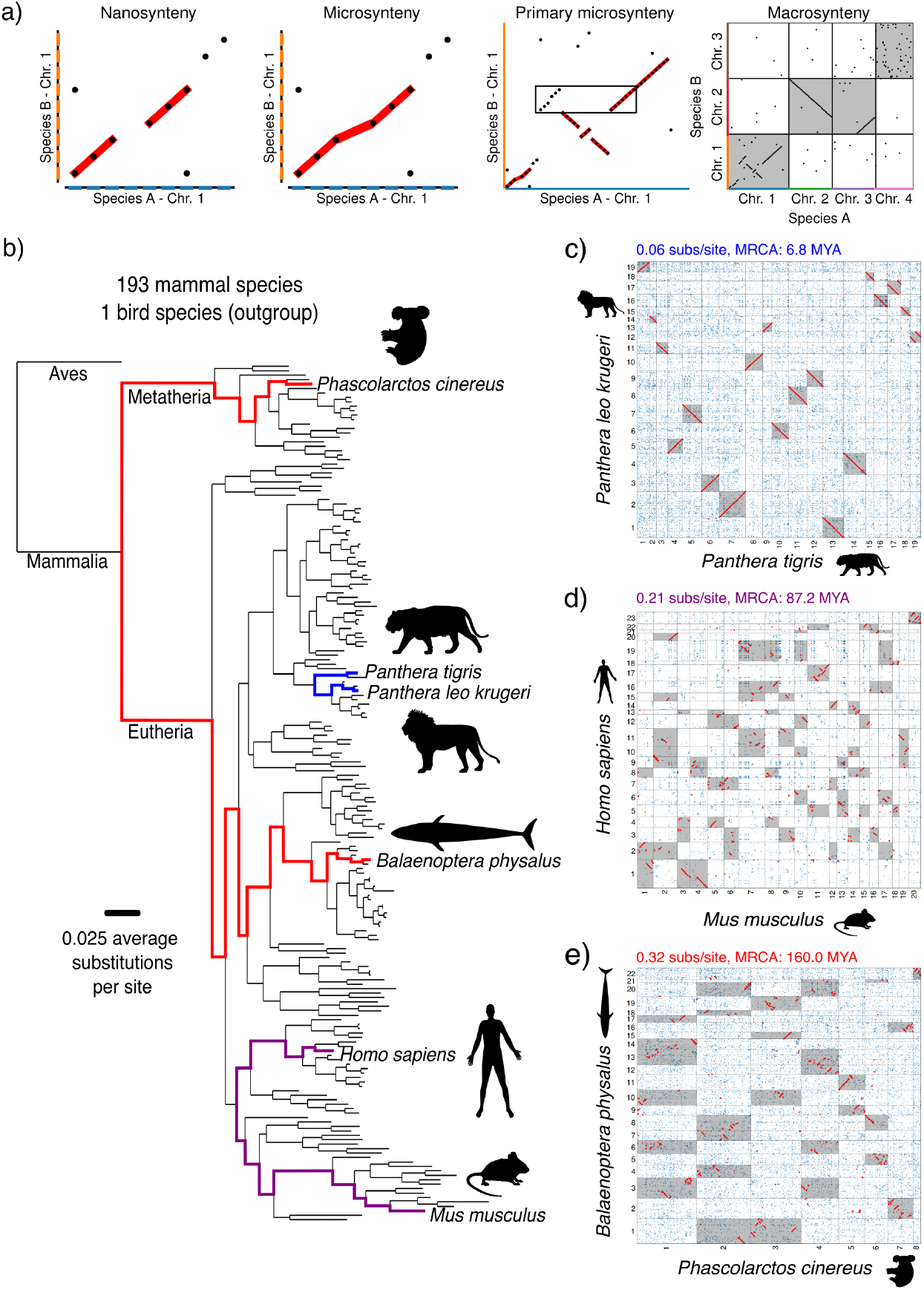
Nanosynteny, microsynteny, and genetic context in mammals. **a)** We define nanosynteny as perfectly preserved gene order, as observed in pairwise dot plots. Microsynteny refers to conserved gene order allowing for small lineage-specific duplications, deletions, translocations, and inversions. Both nanosynteny and microsynteny fit under the widely used definition of collinearity. Primary microsynteny is the largest microsynteny block containing a gene from one of the two genomes in a dot plot. This means that primary microsynteny is defined with respect to a focal and reference species. For example, if we choose species A as the focal species and species B as the reference species, we can see a multi-gene duplication has occurred in the lineage specific to species A. The genes in species B that are syntenic to the two duplicated regions of species B (shown within a box) are contained in a larger synteny block (highlighted in red) and a smaller block. The larger synteny block, primary microsynteny, is likely the ancestral region that was duplicated, and is thus the original context of the duplicated genes before duplication. To distinguish nanosynteny and microsynteny from the traditional meaning of synteny, we use the term macrosynteny, meaning the conservation of genes in microsynteny between two chromosomes. **b)** The phylogeny of the 193 mammalian genome assemblies and single bird outgroup in our dataset, estimated with STAG [52] and STRIDE [53]. Branch lengths are proportional to the amino acid substitutions per site averaged over all orthologs. Three pairwise comparisons are highlighted, corresponding to dot plots between (in order of increasing time to MRCA) **c)** *Panthera leo krugeri* and *Panthera tigris* (0.06 average substitutions per site, MRCA 6.8 MYA) **d)** *Homo sapiens* and *Mus musculus* (0.21 average substitutions per site, MRCA 87.2 MYA) **e)** *Balaenoptera physalus* and *Phascolarctos cinereus* (0.32 average substitutions per site, MRCA 160 MYA). Microsynteny is highlighted in red, and macrosynteny between chromosomes is highlighted in grey. Note the decreasing lengths of microsyntenic regions with increasing time to MRCA, reflecting increased relative fragmentation of genomes with time.

To find microsynteny in mammals, we must first infer homology relationships between genes and then identify conserved gene ordering. We collected chromosome-length genome assemblies from the DNA Zoo [45] and Gencode [46] to assemble a dataset of 193 mammalian genome assemblies and a single bird outgroup (Fig. 1b). While past work on single-gene duplications has relied on a small number of genomes from model organisms, we perform a comprehensive test of location-dependent evolutionary asymmetry in a dense set of mammals. We used OrthoFinder 2 [47, 48] to identify orthogroups (sets of homologous genes across any number of species in the analysis) from translated amino acid sequences (Fig. S1). In order to remove the effects of simple tandem duplications on our results, we condensed each sequence of consecutive genes all in the same orthogroup into a single representative gene, which we also call tandem duplication equivalence classes (TDECs). This commonly used strategy [49, 50, 51] simplifies the identification of multi-gene duplications while retaining information about simple tandem duplications (where a single gene has been duplicated one or more times immediately adjacent to the original copy). Unless explicitly stated otherwise, we use “TDEC” and “gene” interchangeably for ease of reading. We then produced homology matrices between all unique pairs of species, including comparisons between each genome and itself, totaling 18,915 comparisons. A homology matrix is a binary matrix *H*_*ij*_ representing a comparison between species A and B, where *i* runs over all genes in species A and *j* in species B. If genes *i* and *j* are homologous, then *H*_*ij*_ = 1, otherwise *H*_*ij*_ = 0. Homology matrices are visualized as so-called dot plots, which allow rapid inspection of gene order conservation (Fig. 1c-e).

Because of the large amount of data, microsynteny blocks must be inferred algorithmically from homology matrices. Although the problem has already been subject to investigation [49, 50], we developed a novel approach to inferring microsynteny tailored to to address the specific objectives of this study. Our algorithm, dubbed APES (**AP**ES **E**xplore **S**ynteny), identifies synteny at the gene level, which is significantly easier than trying to do the same at the level of nucleotide sequences. Additionally, instead of attempting to recover all synteny, APES is designed to yield a low false positive rate. To achieve this goal, APES relies on first identifying blocks of perfectly conserved gene order, which we call nanosynteny, and then assembling nearby syntenic genes to determine microsynteny blocks (Fig. 2). In this way, each microsynteny block contains at least one nanosynteny block. By randomly permuting homology matrices, we show that nanosynteny blocks containing 3 or more pairs of genes (each pair containing a single orthologous gene from each species in the comparison) are highly unlikely to arise by chance (Fig. 2b-c). The strong statistical significance of nanosynteny blocks with 3 or more putative orthologous gene pairs provides a solid foundation for construction of reliable microsynteny blocks. A complete description of the APES algorithm, together with source code, can be found in the Methods section. Using APES, we identified microsynteny blocks in all species comparisons. In homology matrices, two or more microsynteny blocks containing the same sequence of three or more genes reliably indicate a multi-gene duplication. Next, we sought to determine the source and target copies in these duplications. While other methods exist [54], a straightforward approach to identifying the source copy is to look for the copy with the longest stretch of microsynteny (primary microsynteny) with the reference genome.

**Figure 2:**
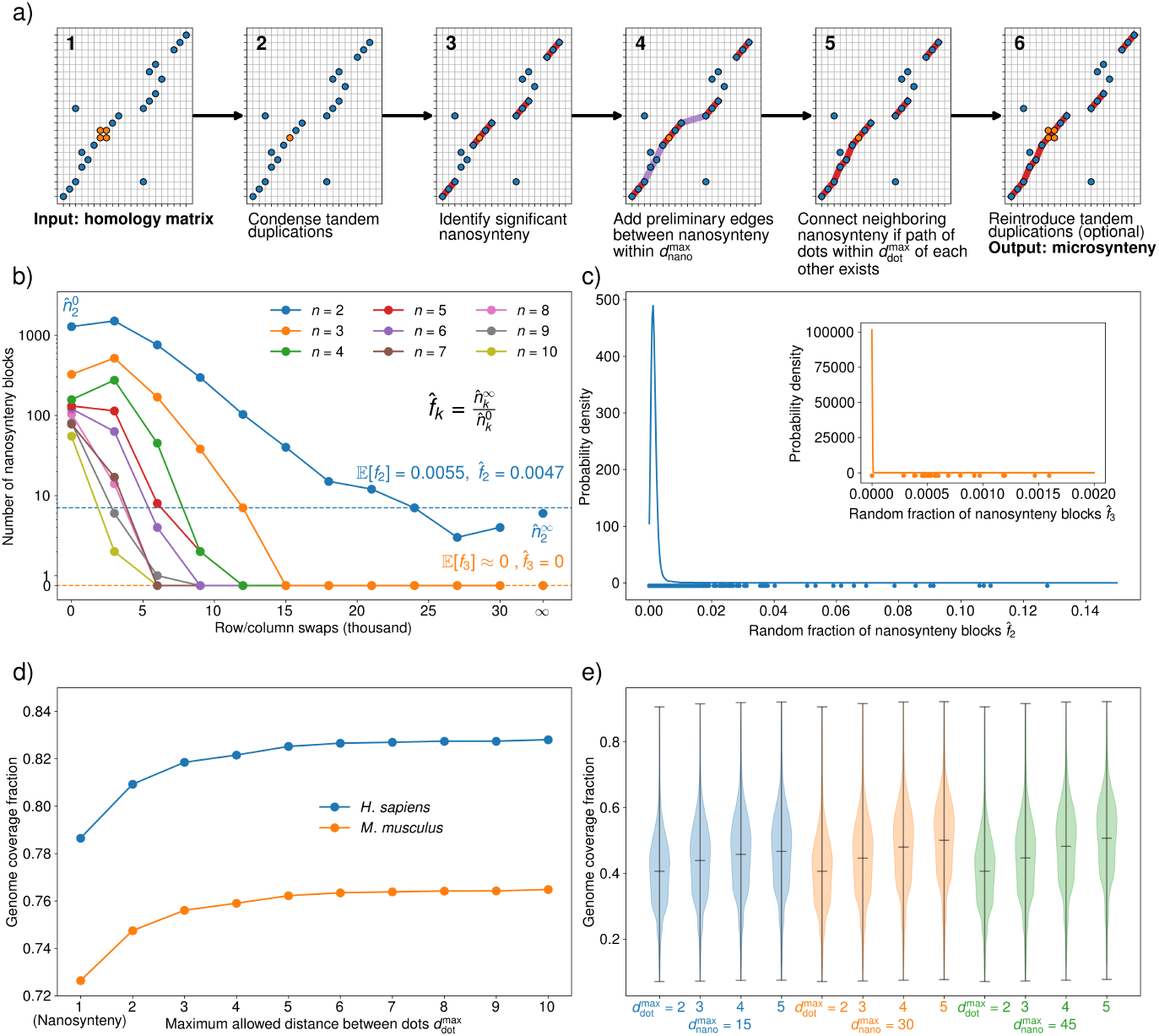
Inferring statistically significant microsynteny. **a)** A schematic of APES for identifying microsynteny blocks, which builds microsynteny blocks from statistically significant nanosynteny blocks in an easily interpretable manner. **Step 1**: The algorithm takes a homology matrix as input, as well as three hyperparameters: the maximum distance allowed between gene Pairs 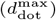, the maximum distance allowed between nanosynteny blocks 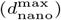, and the minimum anchoring nanosynteny block size. **Step 2**: We find sets of neighboring genes in each genomes which belong to the same orthogroup and condense them into single representative genes for each set, called a tandem duplication equivalence class (TDEC). This prevents false positive microsynteny blocks within large tandem duplication blocks. **Step 3**: Next, we identify all nanosynteny from the homology matrix, retaining nanosynteny blocks containing more than the minimum allowed number of gene pairs. This minimum size parameter can be chosen according to the test detailed in b) and c). **Step 4**: We then add preliminary edges between nanosynteny blocks within 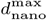 of each other. **Step 5**: For each pair of connected nanosynteny blocks, we check if there is a path of gene pairs within 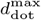 of each other bridging the two nanosynteny blocks. If at least one such path exists, we combine the two nanosynteny blocks into a preliminary microsynteny block along with the shortest identified path of gene pairs. This step is then iterated until all preliminary edges between nanosynteny blocks are checked. **Step 6**: TDECs can optionally then be expanded into their constituent genes and integrated into microsynteny blocks. We do not expand TDECs in this article, so that microsynteny blocks are relationships between TDECs in pairs of genomes. **b)** A permutation test reveals that nanosynteny blocks with more than two gene pairs are unlikely to appear in a homology matrix by chance. We performed random swaps of rows and columns in the *H. sapiens*-*M. musculus* homology matrix, where each swap consisted of randomly choosing and switching two rows and two columns. We then found nanosynteny blocks in the permuted homology matrices. Nanosynteny blocks containing *k* gene pairs are shown in different colored lines for increasing numbers of random row/column swaps. 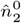 is the number of nanosynteny blocks containing *k* = 2 gene pairs for zero swaps, while 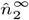 is the same number for 30,000 swaps, where the matrix is effectively random. The number 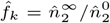 describes the fraction of nanosynteny blocks in the original homology matrix that occur due to random chance. For *k >* 2, the observed fraction is zero. We calculate the expected fractions 𝔼 [*f*_*k*_] from a percolation theoretic statistical model (see Supplemental Information and Fig. S2), finding a close match with the observed fractions. **c)** We performed the analysis in b) for homology matrices between all pairs of species in our dataset, and calculated 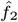 and 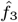 for each. We used kernel density estimates to represent the distributions of 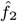 and 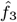 (inset), with the dots below representing the value for each homology matrix. While 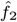 reached up to ∼ 0.25 for some species pairs, 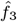 was below 0.003 for all species pairs. We chose a minimum nanosynteny block size of *k* = 3 gene pairs for all subsequent analysis. **d)** Microsynteny blocks are insensitive to maximum allowed distances between gene pairs of 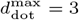 and larger. We identified synteny blocks in the *H. sapiens*-*M. musculus* homology matrix with a range of 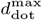 values and measured the fraction of genes in each genome that were within a microsynteny block. By 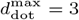, genome coverage fractions for both species plateau, indicating that the genes within microsynteny blocks change little for larger 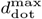values. **e)** We repeated the analysis in d) for all pairs of species in our dataset and used kernel density estimates to represent the distributions of genome coverage fractions. Again, by 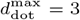 the distribution of genome coverage fractions is similar to the distribution for 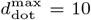, suggesting that the conclusion from d) applies to the dataset as a whole. Varying 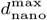 had little effect on the distributions of genome coverage fractions.

We then calculated the Levenshtein distance between each paralog in the source and target copies and the corresponding syntenic ortholog in the reference genome, yielding the number amino acid substitutions per site. Referring to the source paralog as A1, the target paralog as A2, and the syntenic reference ortholog as B1, *d*_*L*_(A1, B1) is the Levenshtein distance between the gene A1 and its syntenic ortholog. The difference Δ*d*_*L*_ = *d*_*L*_(A1, B1) − *d*_*L*_(A2, B1) measures the difference in evolutionary rates between the source and target paralogs with respect to the shared ortholog B1. Δ*d*_*L*_ is negative if the source paralog has experienced a slower substitution rate than the target paralog, positive if the source paralog has experienced a faster substitution rate, and zero if both paralogs have experienced the same substitution rate.

As expected, we found that genes in focal genomes are likely to be most similar to the corresponding reference genome homolog contained in primary microsynteny (Fig. 3). We denote *f* as the fraction of genes in the reference genome for which the homologous focal gene in primary microsynteny is the closest in Levenshtein distance (Δ*d*_*L*_ < 0 for all paralogs) calculated over all cases where Δ*d*_*L*_ ≠ 0. Thus *f* = *P*_*AB*_(Δ*d*_*L*_ < 0|Δ*d*_*L*_ ≠ 0) is defined for an ordered pair of species, where switching species A with species B between the roles of focal and reference species can change *f* . We define *f* ^⋆^ as the same fraction, except calculated only for reference genes belonging to a multi-gene duplication in the focal genome. The fraction *f* ^⋆^ is designed to indicate how often genes in source copies of multi-gene duplications evolve slower than genes in target copies. We find that *f* is large across all species comparisons, with the majority of species comparisons yielding *f* near 0.9 (Fig. 3a-g). Using only nanosynteny yields very similar results (Fig. S3). This result supports the widespread use of synteny in determining orthology. When we restrict our analysis to genes in multi-gene duplications, we find a much broader distribution of *f* ^⋆^ peaked around 0.8 but with very few values less than 0.5 (Fig. 3a-g). While this result seems to suggest asymmetric evolution of copies after multi-gene duplications, a more careful analysis of phylogenetic relationships between focal and reference species is needed. In order to understand the significance of our results, we divided the data into before-and after-speciation duplications, as these two cases are qualitatively different. When the duplication occured before speciation, as in the example shown in Fig. 4a, genes A1 and A2 are paralogs, while A1 and B1 are orthologs, and A2 and B1 are paralogs. This means that the divergence times between A1 and B1 and A2 and B1 are different, obscuring differences in substitution rate. In this case, primary microsynteny does not reliably correlate with the source copy (Fig. 4b). However, when the duplication occurred after speciation as shown in Fig. 4d, differences in substitutions between A1 and B1 compared to A2 and B1 are only due to differences in substitution rate and random chance, not divergence time. Primary microsynteny can be used for after-speciation duplications to identify the source and target copies (Fig. 4e).

**Figure 3:**
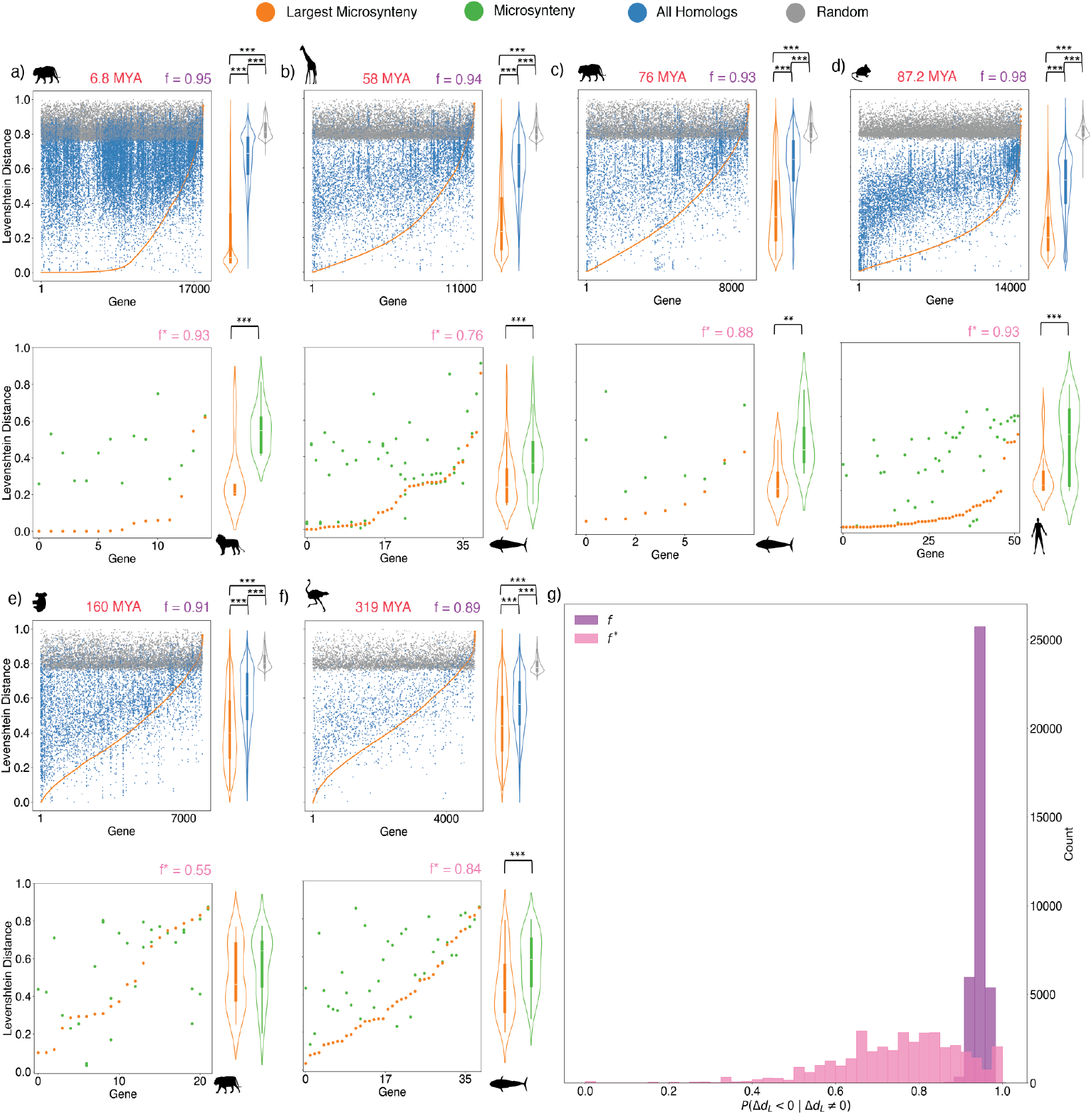
Microsynteny reliably predicts the most similar homolog in pairwise genome comparisons. For each pairwise genome comparison, we designated one species as “reference” and the other as “focal”. We show example pairwise comparisons, written with the reference species first and focal species second, ordered by increasing time to MRCA: **a)** *Panthera leo krugeri* vs. *Panthera tigris*, **b)** *Balaenoptera physalus* vs. *Giraffa camelopardalis*, **c)** *Balaenoptera physalus* vs. *Panthera tigris*, **d)** *Homo sapiens* vs. *Mus musculus*, **e)** *Panthera tigris* vs. *Phascolarctos cinereus*, and **f)** *Balaenoptera physalus* vs. *Struthio camelus*. The upper plots of a)-f) have the indices of all genes with an orthogroup assignment from the reference species shown on the x-axis. Each point on the plot represents comparison of a gene in the reference genome with a homologous (blue and orange) or randomly chosen gene (gray) in the focal species. Orange points with the same x-value represent the unique homolog of a reference species gene among all of its focal species homologs, belonging to the largest microsynteny block. The y-value of each point is the Levenshtein distance between the reference gene and a homolog or randomly chosen gene in the focal species. The value of *f* is the fraction of reference genes for which the corresponding homolog in the largest microsynteny block has the lowest Levenshtein distance of all homologs. This reflects how often a gene in the reference species is least diverged from a focal species homolog in the largest microsynteny block, as compared with other focal species homologs. All distributions of Levenshtein distances for the largest-microsyteny, homolog, and random genes were significantly different from each other (two-sample t-test, *p* < 0.05). The lower plots of a)-f) show similar results, except that only reference genes that belong to at least two microsynteny blocks (i.e. genes duplicated within larger regions) in the comparison with the focal species are shown. *f*^⋆^ is defined similarly to *f* but for the restricted set of genes shown in the lower plots, reflecting how often the larger duplicated region in the focal species is most similar to a given homolog in the reference species. All distributions were significantly different (two-sample t-test, *p* < 0.05), except in e), where *f*^⋆^ = 0.55. **g)** The distributions of *f* and *f*^⋆^ for all 37,442 ordered pairs of species, excluding comparisons of genomes to themselves. The distribution of *f* is sharply peaked around 0.9, demonstrating the utility of synteny in identifying the closest homolog between species. On the other hand, the *f*^⋆^ distribution is far more broad, highlighting the need for an in-depth analysis of duplicated regions.

**Figure 4:**
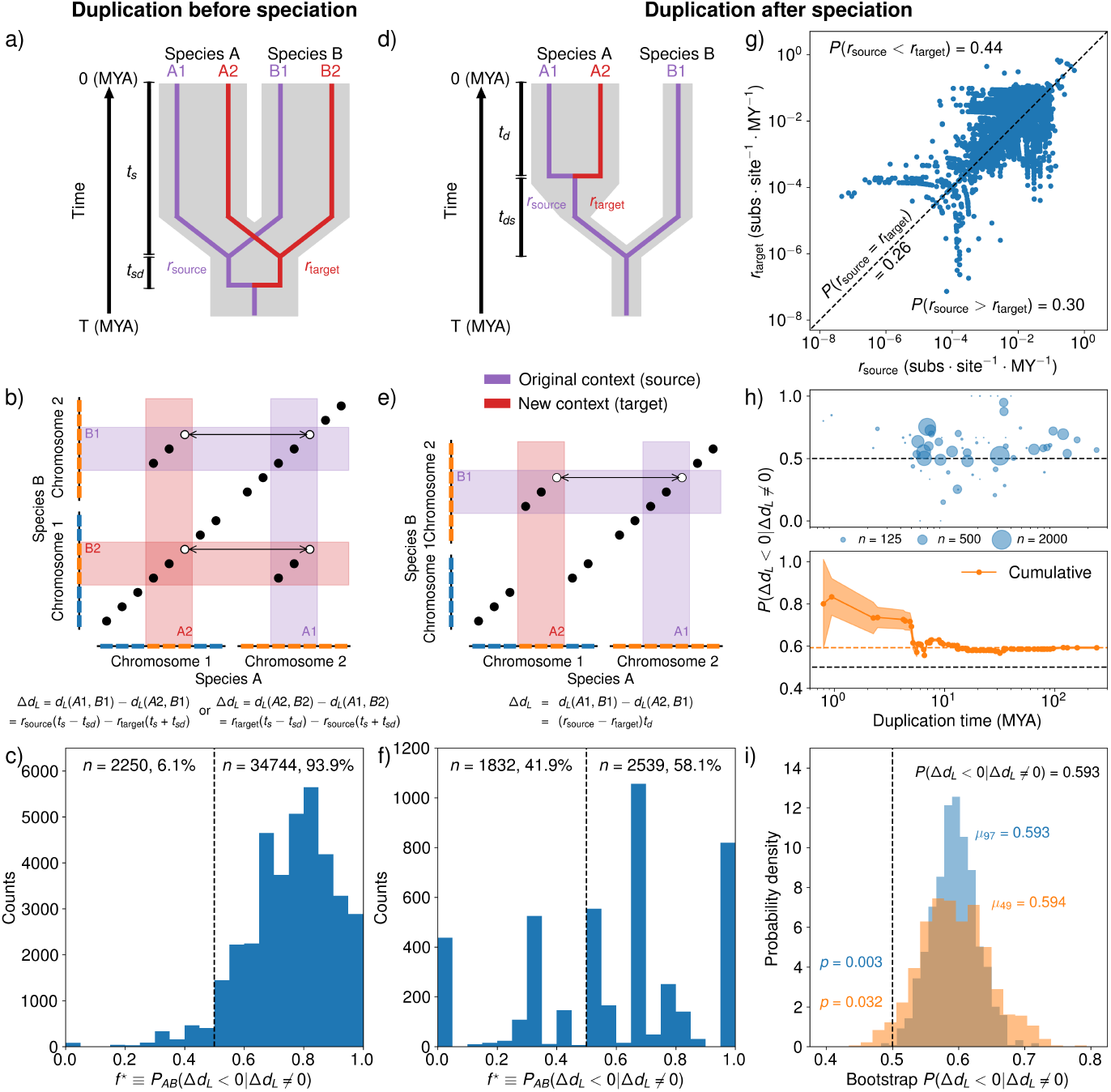
Target gene copies within multi-gene duplications tend to evolve faster source copies. **a)** Illustration of gene duplication before speciation in a two-species comparison. In this example, the two duplicated genes are present in both species A and species B. The large gray tree in the background is the species tree, while the thin tree in the foreground is the gene tree. Species A and species B are separated by a speciation event at *t*_*s*_ million years ago (MYA), which was preceded by a duplication event in the focal orthogroup at *t*_*s*_ + *t*_*sd*_ MYA. The purple branch of the gene tree represents the gene lineage descended from the source copy, while the red branch is the target copy lineage. **b)** A homology matrix (i.e. dot plot) showing evidence of a multi-gene duplication, where each gene within the duplicated region has a gene tree as shown in a). In this case, where the multi-gene duplication occured before speciation, we cannot determine the original context of genes B1 or B2 from primary microsynteny alone. **c)** The distribution of *f*^⋆^ for before-speciation duplications. The *f*^⋆^ distribution is markedly skewed to the right, with 93.9% of *f*^⋆^ values lying above 0.5. This is likely due to gene divergence times, rather than differences in evolutionary rates. Genes A1 and B1 (A2 and B2) in a) and b) diverged more recently than genes A2 and B1 (A1 and B2), so that a greater number of substitutions per site between A2 and B1 (A1 and B2) than between A1 and B1 (A2 and B2) is expected even if *r*_source_ = *r*_target_. **d)** If we instead compare paralogs resulting from a duplication at *t*_*d*_ MYA with an orthologous gene in a species that diverged at *t*_*d*_ + *t*_*ds*_ MYA, before the duplication occurred, we can isolate the effects of differential evolutionary rates. **e)** By calculating the substitutions per site between A1 and B1 and between A2 and B1, we can infer the rate of amino acid substitution. In this case, primary microsynteny is associated with the original context while the smaller microsynteny block contains the target copy of the duplication. **f)** When *f*^⋆^ is calculated for after-speciation comparisons, 58.1% of species pairs exhibit *f*^⋆^ *>* 0.5, indicating that target copies of duplications tend to evolve faster than source copies. **g)** Inferred evolutionary rates for genes in target copies and their paralogs in source copies. For 44% of gene triplets examined, the rate of source copy substitutions was smaller than the rate of target copy substitutions (*r*_source_ < *r*_target_). On the other hand, 30% of gene triplets yielded as larger source substitution rate (*r*_source_ *> r*_target_) and 26% of paralogs evolved at identical rates (*r*_source_ = *r*_target_). Some gene triplets with near-zero *r*_source_ or *r*_target_ are not shown. This demonstrates the asymmetry of paralog evolution after duplication, and the polarization substitution rates so that target copies tend to evolve faster. **h)** For duplications separated by inferred age, no clear trend emerged between rate asymmetry *P* (Δ*d*_*L*_ < 0|Δ*d*_*L*_ ≠ 0) and age (top). We also calculated cumulative *P* (Δ*d*_*L*_ < 0|Δ*d*_*L*_ ≠ 0) for all duplications of age *t* MYA and younger (bottom). Including all gene triplets, we found that 59.3% of gene triplets with non-identical paralogs had an elevated evolutionary rate in the target copy compared with the source copy. **i)** To see how robust our results were to the choice of the species in out data set, we performed bootstrap sampling of species in our analysis without replacement, with either 97 species or 49 species per sample. For both sample sizes, we drew 1,000 samples and calculated *P* (Δ*d*_*L*_ < 0|Δ*d*_*L*_ ≠ 0) using only gene triplets from the sample species. Even with a quarter of the original dataset, the probability of finding *P* (Δ*d*_*L*_ < 0|Δ*d*_*L*_ ≠ 0) ≤ 0.5 remained low (*p* = 0.032). This indicates that our results are not due to the exact set of species we chose to include in our dataset.

We took a conservative approach to estimating the duplication time for each multi-gene duplication through a consensus of two independent methods. Our intent was to minimize the number of times we underestimated the duplication times, to ensure a low false positive rate in determining gene triplets resulting from after-speciation duplications. In the first method, we gathered the duplication times inferred by DLCpar [55] for each paralogous pair in the multi-gene duplication. Within each multi-gene duplication, we then searched for the paralogous pair inferred to arise from a well-supported duplication (50% or more of species that should have inherited the duplication in the absence of deletions have both copies of the gene) with the earliest (largest, in MYA) duplication time. This duplication time was assigned to the entire multi-gene duplication. In the case where none of the paralogous pairs were inferred to be the result of a well-supported duplication, we simply chose the earliest duplication time to represent the multi-gene duplication. Our second method involved searching for the presence of two or more copies of the regions resulting from a multi-gene duplication in each other genome. We then determined the most recent common ancestor of all species in which we found two or more copies of the duplicated regions. The time to this ancestral species should be an upper bound on the duplication time if we assume there cannot be independent multi-gene duplication events of the same region. As a consensus rule, we took the larger of the two calculated duplication times. This process identified multi-gene duplication timings with a fairly uniform distribution (Fig. S4). Because after-speciation duplications directly reflect the asymmetry of evolutionary rates between the source and target copies, we use this information to characterize the effect of genetic context in the evolutionary rate of otherwise identical copies at the time of duplication. For the after-speciation subset, we estimated the substitution rates of the source (*r*_source_) and target (*r*_target_) copies. Of these 22,342 gene triplets, 26% had no difference in rate (*r*_source_ = *r*_target_). Another 30% of gene triplets had a higher substitution rate for source copy (*r*_source_ *> r*_target_) while 44% had a lower substitution rate for source copy (*r*_source_ < *r*_target_) (Fig. 4g). This amounted to a probability of *P* (Δ*d*_*L*_ < 0|Δ*d*_*L*_ ≠ 0) = 0.593 overall, so that roughly 60% of all cases where there is a measurable rate difference between paralogs in multi-gene duplications, the target copy evolves faster than the source copy. All results were similar when we replaced microsynteny analysis with nanosynteny (Fig. S5). Note that the probability *P* (Δ*d*_*L*_ < 0|Δ*d*_*L*_ ≠ 0) pools all species pairs together, in contrast to *P*_*AB*_(Δ*d*_*L*_ < 0|Δ*d*_*L*_ ≠ 0) which is defined with respect to an ordered pair of species.

We did not find a strong relationship between rate asymmetry and duplication time. Visually, *P* (Δ*d*_*L*_ < 0|Δ*d*_*L*_ ≠ 0) had little dependence on duplication time (Fig. 4h), while the correlation between the sign of Δ*d*_*L*_ and duplication time was very weak (*ρ* = −0.042, *p* = 4.35 · 10^−10^). This result is inconsistent with previous findings of accelerated rates in young duplications [13, 15]. However, we did not focus our efforts on estimating fine-grained, branch-specific rates and further work is needed to examine the issue of rate acceleration and deceleration after duplication in mammals.

Our analysis to this point implicitly assumes that each gene triplet is independent from all others. This assumption can fail when two gene triplets result from the same duplication event. This can occur in two distinct ways. First, two gene triplets can differ only by the reference ortholog. Second, one gene triplet can contain a pair of paralogs from one species while another gene triplet contains a distinct pair of paralogs from another species. Both pairs of paralogs can derive from a single duplication event followed by a speciation event.

We used several approaches to address the impact of dependent gene triplets on our results. First, we performed bootstrap subsampling of the species in our analysis, effectively reducing the impact of dependence by decreasing the probability of choosing two species that share two or more copies of a genomic region resulting from a duplication. Bootstrap subsampling yielded a low probability of finding *P* (Δ*d*_*L*_ < 0|Δ*d*_*L*_ ≠ 0) < 0.5 even when using only a quarter of the species from our dataset (Fig. 4i). This shows that our results are robust to significant reductions in the dataset, suggesting the apparent polarization in rate asymmetry is not a consequence of either dependent data or the specific choice of species for analysis. We also performed our analysis after averaging over all possible reference orthologs for each pair of paralogs and separately for only pairs of paralogs resulting from lineage-specific duplications. Averaging over reference orthologs removes the possibility of comparing two gene triplets that are identical except for the reference ortholog, while examining terminal duplications averaged over reference orthologs fully ensures that each gene triplet is independent. Additionally, using a null model of amino acid substitution to account for sequence differences that can occur by random chance even when *r*_source_ = *r*_target_, we find strong polarization for significantly different paralog sequences, for both non-terminal and terminal duplications (Fig. S6). Next, we sought to examine how the differences in the regulatory environments of the source and target copies impact the polarity of asymmetric evolution. We expect that the larger that a duplicated region is, the greater the chance is that the regulatory environment around a gene is carried to the target location in the genome. Consequently, we expected to see polarization decrease with the size of the duplication if asymmetry in regulatory environment between the source and target copies contributes to polarization. Accordingly, we found that polarity decreased with larger target copies, suggesting that larger duplications lead to lower polarity (Fig. 5a). For small duplications, i.e. when the target region is small, the substitution rate *r*_target_ is far more likely to be elevated over *r*_source_ than for large duplications. Notably, in the largest class of duplications, we still found polarization, suggesting that regulatory environments can extend past ∼ 1 Mb or that other factors influence polarity. Such long-range regulatory activity is consistent with the very largest enhancer-to-transcription start site distances found in the human genome [56]. Overall, the inverse relationship we found between polarization and duplication size supports the hypothesis that larger duplications are more likely to carry cis-regulatory elements and important co-expressed genes to the target genomic location, creating a similar regulatory environment to the source location and increasing purifying selection on target copy genes.

**Figure 5:**
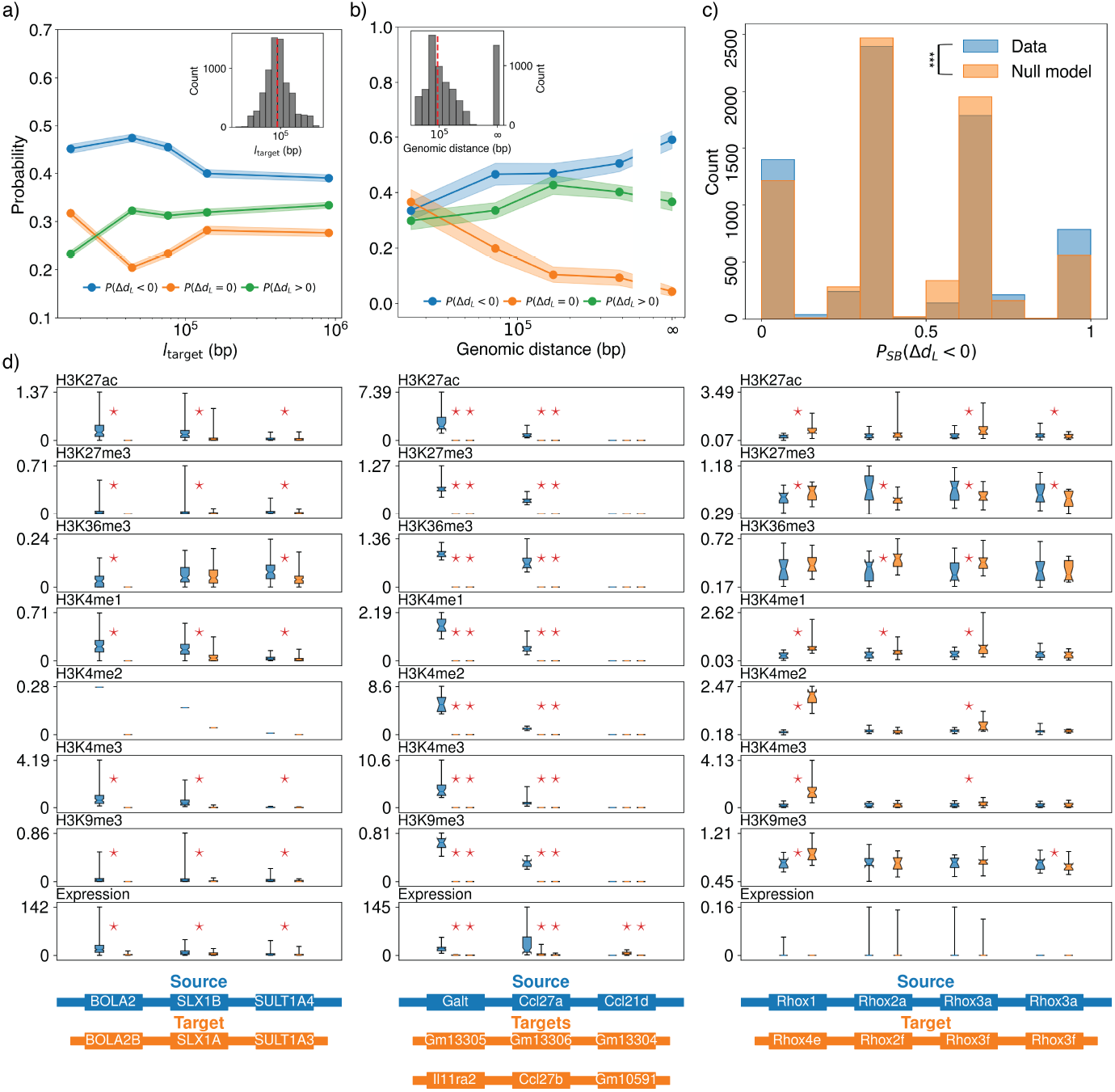
The fate of duplicated genes is influenced by the size of the duplicated region and the genomic distance between copies, and is shared among the genes within a multi-genic copy. **a)** The probability of the source copy evolving slower than the target copy, *P* (Δ*d*_*L*_ < 0), falls with larger target copy size in basepairs, *l*_target_, while the probability of the source copy evolving faster, *P* (Δ*d*_*L*_ *>* 0), increases. The difference between the two probabilities represents the polarity of asymmetric paralog evolution, such that when *P* (Δ*d*_*L*_ < 0) < *P* (Δ*d*_*L*_ *>* 0), the source copy is more likely to be the slowly evolving copy. Inset: the distribution of target copy sizes resulting from multi-gene duplications. **b)** When the same probabilities are examined as a function of the genomic distance between copies, there is no obvious evidence of whether the polarity of asymmetric evolution is influenced by the genomic distances inter-spacing the two copies. In this case both *P* (Δ*d*_*L*_ < 0) and *P* (Δ*d*_*L*_ *>* 0) increase with increased genomic distance, and we observe a monotonic decrease in *P* (Δ*d*_*L*_ = 0) with genomic distance. Nonetheless, remarkably, when the two copies are on different chromosomes (genomic distance of ∞), the difference between *P* (Δ*d*_*L*_ < 0) and *P* (Δ*d*_*L*_ *>* 0) is greatest, and polarization is maximal. **c)** The probability of the source copy evolving slower than the target copy within a multi-gene duplication (*P*_*SB*_ (Δ*d*_*L*_), *SB* for synteny block) is significantly different from a null model produced by randomly resorting paralogous genes into new multi-gene duplications. If all paralogous gene pairs within a large duplicated region were polarized in evolutionary rate towards either the target or source copy, we would expect to see only values of *P*_*SB*_ (Δ*d*_*L*_) = 0 or *P*_*SB*_ (Δ*d*_*L*_) = 1. When compared with an ensemble of gene triplets randomly assigned to multi-gene duplications, the probability of observing our original data was low (*p* < 0.05), with excess counts at *P*_*SB*_ (Δ*d*_*L*_) = 0 and *P*_*SB*_ (Δ*d*_*L*_) = 1. This suggests that the fate of multi-gene duplications is determined collectively more often than expected by random chance. All results in a)-c) were similar when considering microsynteny or only nanosynteny (Figs. S7 & S8). **d)** Analysis of ChIP-seq data for epigenetic markings associated with expression and RNA-seq data suggests that both may correlate with the polarity of asymmetric evolution in multi-gene duplications. While, only three sets of regions in the human and mouse genomes satisfied our criteria for analysis of ChIP-seq and RNA-seq data (i.e. that we could unambiguously distinguish the source and target regions), we found significant differences (Wilcoxon signed-rank test, *p* < 0.05) across most paralogs in both epigenetic markings and expression for the two sets of regions on autosomal chromosomes in human (first column) and mouse (second column). In the last case, a large duplicated region in the mouse X chromosome (third column), there were fewer significant differences for epigenetic markings and no significant differences in expression. Note that in this analysis we examine individual genes, rather than TDECs.

We also expected polarity to increase with the genomic distance between copies, reasoning that distant target copies would experience different regulatory environments from the source copy while nearby target copies would experience similar regulatory environments. The dependence of polarization on genomic distance between the two copies is less clear, with polarity appearing more or less insensitive for intrachromosomal duplications (Fig. 5b). However, polarity was maximal for copies on different chromosomes, consistent with previously reported results in *C. elegens* [12]. While increasing distance between copies on the same chromosome appears to have a moderate influence on polarity, the novel regulatory environment faced by target copies on different chromosomes from their source copies may explain the polarity found in their evolutionary rates.

While paralogous pairs within large duplicated regions clearly demonstrate polarity, we sought to determine whether paralogous pairs created simultaneously in a single multi-gene duplication event share polarities. Since the genes within a single duplicated region share a common regulatory environment, we expect pairs of paralogs created in a single multi-gene duplication to be have correlated polarities, so that the “choice” of the slower and faster evolving paralog occurs in a block-wise manner. To test this expectation, we defined the within-synteny block probability *P*_*SB*_(Δ*d*_*L*_ < 0) as the fraction of gene triplets within a duplicated region with Δ*d*_*L*_ < 0. If all genes on in the source copy have evolved at a slower rate than their paralogs on the target copy *P*_*SB*_(Δ*d*_*L*_ < 0) will be one, while *P*_*SB*_(Δ*d*_*L*_ < 0) will be zero if the source copy genes have evolved faster than, or at the same as, their target copy paralogs. Intermediate probabilities reflect mixed polarities within duplicated regions. Using a null model that randomly assigns gene triplets to empirically determined duplicated regions, simulating what we would expect if the polarity of paralogous gene pairs was independently determined, we found that the *P*_*SB*_(Δ*d*_*L*_ < 0) distribution displayed significant enrichment of *P*_*SB*_(Δ*d*_*L*_ < 0) = 0 and *P*_*SB*_(Δ*d*_*L*_ < 0) = 1. This is strong evidence that polarity determination in multi-gene duplica-tions tends to occur in a block-wise manner rather than independently for each paralogous gene pair, suggesting that the fate of a duplicated gene can be influenced by the other genes and non-coding elements simultaneously duplicated.

Finally, we explored the role of differential epigenetic markings and gene expression on asymmetric evolution of paralogs. We restricted our analysis of epigenetic markings to histone modifications with known associations with gene expression [57, 58, 59, 60] (Table S1). Unfortunately, experiments probing histone modifications are routinely performed only on a handful of organisms; out of the 194 species in our data set, experimental data was available only for humans and mice. Out of all multi-gene duplications in human and mouse, only three could be reliably placed in the after-speciation category. According to our analysis, the human genome presents a single multi-gene duplication after speciation, which shows significant differences in both epigenetic markings and expression levels between the source and target copies, with higher markings and expression levels in the source copy (Fig. 5d). The distributions of expression levels and epigenetic marks represented in Fig. 5d are for all cell types for which data were available. Thus, a broad distribution (with a wide range of expression or epigenetic marking levels) indicates variability across tissues, while a narrow distribution corresponds to homogeneity across tissues. The first set of duplicated mouse regions had two target copies associated with the source copy, and similarly had significant differences between the source copy and both targets. The second multi-gene mouse duplication was on the X chromosome, in contrast to the previous two duplications which had both copies on autosomal chromosomes. In this duplication, the source and target copies had fewer significant differences in epigenetic markings and gene expression was not significantly different between any pair of paralogs. Overall, these results are consistent with some role of epigenetic markings in polarization and correlate higher expression levels with slower substitution rates, as expected.

In conclusion, our findings provide substantial evidence for asymmetry and polarization in the amino acid substitution rate of the two copies (source and target) resulting from a multi-gene duplications in mammals. Our results are particularly significant in virtue of the large dataset we used. In short, our analysis indicates an important role for location in polarizing substitution rates of paralogs: the source copy in the ancestral genomic location tends to evolve slower than the target copy in the new locus created by the duplication event. By focusing on multi-gene duplications, we were able to demonstrate that polarization decreases with the size of the duplicated region, yet remains significant for duplications up to the scale of at least one megabase. Moreover, we demonstrate that the asymmetry in substitution rate tends to occur in a block-wise manner, so that genes in one copy tend to all evolve slower or faster than their paralogs in the other copy.

The statistical methods we used in synteny detection are central to our approach. Leveraging the strict definition of nanosynteny, we developed a mathematically rigorous estimate of statistical significance for nanosynteny blocks. Using nanosintheny blocks as building blocks for microsynteny, we were then able to identify reliable regions of conserved gene order across a large, representative set of mammals. APES allowed for easy interpretation of synteny blocks, where commonly used dynamic programming algorithms can yield conflicting synteny blocks without any notion of statistical significance. Overall, this innovative approach brings together several previously isolated observations regarding the evolutionary rates of duplicated genes, offering a quantitative analysis of how genetic context influences duplicated genes in mammals.

## Methods

### Genome assemblies and annotations

We downloaded MAKER2 [61] genome annotations of 191 mammals and 1 bird (as an outgroup) from the DNA Zoo [45], along with the GENCODE human genome annotation v43 and mouse genome annotation v32 [46].

### Orthogroup detection

We used OrthoFinder 2 [47, 48] to determine orthogroups, using DIAMOND [62] to identify putative homology relationships between genes. DIAMOND alignment scores are used as the edges of an orthogroup graph, with genes as nodes. The Markov Cluster Algorithm (MCL) [63] is then used to find communities in the orthogroup graph, yielding disjoint orthogroups. OrthoFinder also constructs a rooted species tree [53, 52] and rooted gene trees for each orthogroup, reconciled with the species tree through DLCpar [55]. DLCpar finds parsimonious gene tree reconciliations considering gene duplication, gene loss, and deep coalescence.

### Microsynteny inference with APES

We developed an algorithm called APES (**AP**ES **E**xplore **S**ynteny) in order to reliably infer microsynteny with a low chance of yielding false positive results. APES infers microsynteny in each pair of species by first finding regions of perfectly conserved gene order (nanosynteny), and subsequently piecing together statistically significant nanosynteny into microsynteny. Nanosynteny blocks were considered statistically significant if they contained three or more TDEC pairs in order to ensure that the expected fraction of nanosynteny blocks found purely due to chance were less than 0.05 for all genome comparisons (Fig. 2). In order to avoid including regions only representing more complex tandem duplications, we filtered out nanosynteny blocks containing fewer than three unique orthogroups. We then joined nanosynteny blocks based on two hyperparameter choices, controlling the maximum distance between nanosynteny blocks in both genomes, 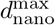, and the maximum distance of gene pairs from each other in the path between the nanosynteny blocks, 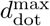. See Fig. 2 and the Supplemental Information for details. Python code is available at https://github.com/DiPierroLab/GenomicTools.

### Finding multi-gene duplications and associated timings

We identified multi-gene duplications in each genome using within-genome microsynteny. We applied a series of filtering steps to the within-genome microsynteny blocks as follows:

1. We first excluded all microsynteny blocks representing alignment of each gene in a chromosome with itself.
2. We next removed any remaining microsynteny blocks that contain the same gene twice.
3. Finally, we removed overlaps between microsynteny blocks. If any two microsynteny blocks were on the same two chromosomes, the overlap is between them is defined as the smaller number of shared genes in both blocks between the first and second genome (see Supplemental Information for details). The larger (measured in number of TDECs) of two overlapping blocks was retained, while the smaller block was removed.

All analysis of multi-gene duplications was performed on the remaining pairs of microsyntenic regions in each genome. We remained intentionally agnostic as to the mechanism producing large duplicate regions, though these could be a result of segmental duplications, whole-genome duplications with subsequent chromosomal rearrangements, or even complex tandem duplications.

In order to infer the time of each multi-gene duplication event, we used the species tree and reconciled gene trees produced by OrthoFinder. Because OrthoFinder does not take synteny into account, the reconciliation process is independent for each orthogroup. However, in multi-gene duplications we must introduce the constraint that all paralogous pairs were produced simultaneously in a single duplication event. While reconciliation methods exist for this problem [64, 65], we took a simpler approach designed to reduce the probability of underestimating duplication times (measured in millions of years from the present). We used a consensus of two independent methods. First, we used the reconciled gene trees from OrthoFinder. Each paralogous pair is assigned to a duplication event at a node in the species tree. We will refer to this as the duplication node. Each duplication node has an associated support, defined as the fraction of descendant leaves (i.e. extant animal genomes in the dataset) of the duplication node that have retained both paralogs resulting from the duplication event. We defined the duplication node of a multi-gene duplication as the earliest (closest to the root node), well-supported (≥ 0.5) duplication node out of all the paralogous pairs therein. If none were well-supported, then we simply took the earliest duplication node. Second, we searched for the sequence of orthogroups found in the multi-gene duplication in all genomes. If two or more copies of those orthogroup sequences were found in a species we added it to a list of multiple-copy species. The duplication node was then simply the MRCA of all multiple-copy species in the species tree.

We then took the earlier of the two duplication nodes from our two methods as the consensus duplication node. Using divergence times from TimeTree 5 [66], the duplication time is defined as the time halfway between the divergence time associated with the duplication node and its parent species tree node.

### Substitution rate inference

We estimate amino acid substitution rates, rather than estimating synonymous and non-synonymous nucleotide substitution rates. Using amino acid substitution rates is justified in our analysis because of the large divergence times between the species in our dataset (such that non-synonymous nucleotide substitutions are in danger of saturation), and because we have reasonable estimates of divergence times between paralogs from OrthoFinder 2 and TimeTree. We calculate substitution rates by dividing Levenshtein distances by duplication times. See the Supplemental Information for details.

### Epigenetics and gene expression data

We downloaded human and mouse histone ChIP-seq data from the ENCODE project [67, 68, 69] for the markings in Table S1. For each marking, we calculated the average fold change over control signal over each relevant gene for each tissue type available. This left us with an ensemble of marking signals for each human and mouse gene. We downloaded human and mouse gene expression data for the tissues listed below from the ENCODE project using RNAget [70].

## Acknowledgments

We thank the members of the Nuclear Physics working group in the Center for Theoretical Biological Physics for valuable discussion. We are especially grateful to Erez Aiden and Olga Dudchenko for their incisive input. This research was supported by the National Science Foundation through the awards PHY-2412651 and PHY-2019745, and by the National Institute of General Medical Sciences of the NIH under award R35-GM146852. The content is solely the responsibility of the authors and does not necessarily represent the official views of the funding agencies.

## Notes

https://github.com/DiPierroLab/GenomicTools

### Competing Interest Statement

The authors have declared no competing interest.

### Summary of Updates

Updates to text for clarity, Figure 1 revised.

