## Supplemental Information for "Quantifying the influence of genetic context on duplicated mammalian genes"

### Supporting information for: Quantifying the influence of genetic context on duplicated mammalian genes

May 2, 2025

### 1 Genome annotations

#### 1.1 Notation

We denote the set of species in our analysis as $\mathcal{S}$. A species $S \in \mathcal{S}$, has a set of chromosomes, represented by the index set $\mathcal{C}^S = \{1, 2, \ldots, C^S\}$ where $C^S$ is the haploid number of chromosomes in species $S$. The index of a chromosome $c \in \mathcal{C}^S$ corresponds to the usual numbering of chromosomes. We consider only autosomal and X chromosomes, as the genomes in our dataset are sampled from a mixture of male and female individual mammals.

We write the genes of species $S$ in chromosome $c$ as the ordered set

$$G_c^S = \{g_{c1}^S, g_{c2}^S, \ldots, g_{cN_c^S}^S\} \tag{1}$$

where $N_c^S$ is the number of genes on chromosome $c$ of species $S$. The indices represent the order of the genes on the chromosome (with the $i = 1$ side of the chromosome chosen arbitrarily) so that

$$g_{ci}^S < g_{cj}^S \iff i < j. \tag{2}$$

The set of all genes in a species

$$G^S = \bigcup_{c \in \mathcal{C}} G_c^S \tag{3}$$

is partially ordered according to the gene order within each chromosome (meaning that there is no order relationship between genes from different chromosomes).

##### 1.2 Data

We downloaded genome annotations of 191 mammals and 1 bird (as an outgroup) from the DNA Zoo [1], along with the GENCODE human genome annotation v43 and mouse genome annotation v32 [2].

#### 2 Inferring homology of protein-coding genes

##### 2.1 Notation

We write the set of all orthogroups assigned by OrthoFinder as

$$\mathcal{O} = \{0, 1, \ldots, O\}, \tag{4}$$

where $O$ is the total number of orthogroups found; all genes for which OrthoFinder could not assign an orthogroup were assigned to the orthogroup index 0. We denote the orthogroups of genes through the function

$$o : \bigcup_{S \in \mathcal{S}} G^S \to \mathcal{O}. \tag{5}$$

This function maps the set of all genes across all species and chromosomes to orthogroup indices.

##### 2.2 Running OrthoFinder

We used OrthoFinder 2 [3, 4] to find the orthogroups defined by our set of genomes from 194 species, $\mathcal{S}$. For each genome, we took the amino acid sequences of the largest isoform associated with each protein-coding gene. Annotations of the human and mouse genomes were downloaded from GENCODE, while the remaining annotations were downloaded from the DNA Zoo and were created with MAKER 2 [5]. OrthoFinder uses DIAMOND [6] to infer homology between genes, correcting scores for amino acid sequence length and phylogenetic distance. Genes are then assembled into a graph with weighted edges reflecting thresholded, corrected DIAMOND scores and the Markov cluster algorithm (MCL) [7] is used to find orthogroups. We used an inflation parameter of 1.5 in MCL.

OrthoFinder 2 [4] additionally infers a rooted species tree [8, 9] and gene trees for each orthogroup reconciled to the species tree through the DLCpar algorithm [10]. As the STRIDE algorithm incorrectly placed the root of the species tree, we chose an alternate species tree provided by STRIDE with the root correctly placing the single non-mammal (ostrich) as an outgroup. The inferred species has branch lengths reflecting the substitutions per site averaged over all protein-coding genes.

### 3 Condensing tandem duplications

We follow the common practice of condensing tandem duplications before identifying synteny blocks [11, 12, 13]. This process effectively removes the simplest type of tandem duplication detectable at the gene scale, where a gene is copied and both copies are on the chromosome with no other genes between them. We define a tandem duplication block as a sequential set of genes on a chromosome all belonging to the same orthogroup. Formally, we define tandem duplication blocks through the equivalence relation $\sim$ between two genes on chromosome $c$, $g^S_{ci}$ and $g^S_{cj}$, according to

$$g^S_{ci} \sim g^S_{cj} \iff o(g^S_{ci}) = o(g^S_{cj}) \neq 0 \text{ and } |i - j| \leq 1. \tag{6}$$

This equivalence relation generates equivalence classes

$$\tilde{G}^S_c = G^S_c / \sim \tag{7}$$

which partition $G^S_c$ into tandem duplication blocks. We interchangeably call these equivalence classes condensed genes and tandem duplication equivalence classes (TDECs). Note that a TDEC can contain a single gene, which is a “trivial” tandem duplication only in a formal sense.

The ordering of $\tilde{G}^S_c$ is inherited from $G^S_c$, meaning that for $\tilde{g}^S_{ci}, \tilde{g}^S_{cj} \in \tilde{G}^S_c$

$$\tilde{g}^S_{ci} < \tilde{g}^S_{cj} \iff g^S_{ck} < g^S_{cl} \text{ for some } g^S_{ck} \in \tilde{g}^S_{ci} \text{ and } g^S_{cl} \in \tilde{g}^S_{cj}. \tag{8}$$

TDECs are indexed for a given species and chromosome starting at $i = 1$ for the TDEC containing $g^S_{c1}$, $i = 2$ for the TDEC containing $g^S_{ck}$ where $k$ is the smallest gene index such that $g^S_{ck}$ is not in $\tilde{g}^S_{c1}$, and so on.

Abusing notation, we use the orthogroup $o$ function to denote the orthogroup of a TDEC

$$o(\tilde{g}^S_{ci}) = o(g) \text{ for all } g \in \tilde{g}^S_{ci}. \tag{9}$$

### 4 Inferring microsynteny using APES

We designed a conservative microsynteny algorithm called APES (**AP**ES **E**xplore **S**ynteny) that eschews scoring functions and dynamic programming. The algorithm first identifies perfect gene order conservation, which we call nanosynteny, in a condensed homology matrix comparing two genomes, and then connects nanosynteny blocks into microsynteny blocks according to a simple rule.

#### 4.1 Parameters

APES requires three parameter choices. The first parameter is the minimum number of genes required for a nanosynteny block to be retained, $k_{\min}$. Requiring that nanosynteny blocks have at least a minimum number of genes in them avoids the possibility of spurious nanosynteny blocks occurring due to permissive gene homology inference. As we will show below, we can directly estimate the appropriate value of this parameter from a condensed homology matrix.

The second parameter is the maximum distance allowed between nanosynteny blocks for them to be candidates for connection into a microsynteny block, $d_{\text{nano}}^{\max}$. We use the following metric to measure distance between "dots", meaning pairs of homologous condensed genes, one from each species being compared, in the condensed homology matrix

$$d_{\mathrm{H}}(\mathbf{x}, \mathbf{x}') = \max\left(|i-k|, |j-l|\right) \tag{10}$$

where $\mathbf{x} = (\tilde{g}^A_{ci}, \tilde{g}^B_{c'j})$ and $\mathbf{x}' = (\tilde{g}^A_{ck}, \tilde{g}^B_{c'l})$ are the pairs of homologous condensed genes in chromosome $c$ of species $A$ and chromosome $c'$ of species $B$. Two nanosynteny blocks are only allowed to be connected if they are on the same pair of chromosomes and have the same slope. For a nanosynteny block $\mathbf{s} = \{\mathbf{x}_1, \mathbf{x}_2, \ldots, \mathbf{x}_n\}$, we define the slope as

$$\text{slope}(\mathbf{s}) = \text{sign}\left(\frac{l-j}{k-i}\right) \tag{11}$$

for $\mathbf{x}_1 = (\tilde{g}^A_{ci}, \tilde{g}^B_{c'j})$ and $\mathbf{x}_n = (\tilde{g}^A_{ck}, \tilde{g}^B_{c'l})$. Note that the slope of each individual nanosynteny block is arbitrary, depending on which end of each chromosome in each genome is picked to have an index of 1. However, once indexing decisions have been made for each genome, two nanosynteny blocks of the slope potentially represent a larger region of imperfectly conserved gene order. The distance between two nanosynteny blocks with the same slope is

$$d_{\text{nano}}(\mathbf{s}, \mathbf{s}') = \min\left(\{d_H(\mathbf{x}, \mathbf{x}') \mid \mathbf{x} \in \mathbf{s}, \mathbf{x}' \in \mathbf{s}'\}\right). \tag{12}$$

Two nanosynteny blocks with the same slope and on the same pair of chromosomes are then candidates for connection if

$$d_{\text{nano}}(\mathbf{s}, \mathbf{s}') \leq d_{\text{nano}}^{\max}. \tag{13}$$

The final parameter is the maximum distance allowed between dots, $d_{\text{dot}}^{\max}$, used in deciding whether two candidate nanosynteny blocks should be connected. We use the metric in Eq. 10 between pairs of dots, and allow for a connection between candidate nanosynteny blocks if there is a path between them of dots each within $d_{\text{dot}}^{\max}$ of each other. See the full description of the algorithm below for more details.

In order to settle on reasonable values of the $d_{\text{nano}}^{\max}$ and $d_{\text{dot}}^{\max}$ parameters, we repeated our analysis with varied values of both. We chose values of $d_{\text{nano}}^{\max}$ and $d_{\text{dot}}^{\max}$ in regions of parameter space where the coverage of genomes by microsynteny changed little with small changes in the parameters.

##### 4.2 Step 1: Condense tandem duplications

In the first step, we condense each genome so that tandem duplications are considered as a single condensed gene, as described in Section 3.

##### 4.3 Step 2: Find nanosynteny blocks

We next identify nanosynteny blocks from the condensed homology matrix. A condensed homology matrix for two species $a$ and $b$ is defined as

$$D^{AB}_{cc'ij} = \begin{cases} 1 & o(\tilde{g}^A_{ci}) = o(\tilde{g}^B_{c'j}) \\ 0 & o(\tilde{g}^A_{ci}) \neq o(\tilde{g}^B_{c'j}) \end{cases} \tag{14}$$

where $\tilde{g}^A_{ci}$ and $\tilde{g}^A_{ci}$ are condensed genes within chromosomes $c$ and $c'$ of species $A$ and $B$, respectively. We identify nanosynteny blocks from the condensed homology matrix by simply looking for sequences of condensed dots with sequential indices in both organisms, with either increasing or decreasing indices. This represents perfect locally conserved gene order.

The total number of genes in chromosome $c$ of species $S$ is $\tilde{N}^S_c = |\tilde{G}^S_c|$ while the total number of genes in species $s$ is

$$\tilde{N}^S = \sum_{c=1}^{C^S} \tilde{N}^S_c. \tag{15}$$

We can estimate the probability of observing $n$ nanosynteny blocks of size $k$ under randomization of gene order in both species, using the density of dots

$$\rho = \frac{1}{\tilde{N}^A \tilde{N}^B} \sum_{c=1}^{C^A} \sum_{c'=1}^{C^B} \sum_{i=1}^{\tilde{N}^A_c} \sum_{j=1}^{\tilde{N}^B_{c'}} D^{AB}_{cc'ij}. \tag{16}$$

In accordance with a basic result from percolation theory, the probability that we will observe exactly $k$ dots in a perfect diagonal by random chance is

$$r_k = \rho^k (1-\rho)^2. \tag{17}$$

With this probability in hand, and considering that nanosynteny blocks can occur with positive or negative slope and can be located anywhere in the homology matrix, the probability of observing $n$ synteny blocks of size $k$ by random chance is approximately

$$p_k(n) = \binom{2\tilde{N}_a \tilde{N}_b}{n} (1 - r_k)^{2\tilde{N}_a \tilde{N}_b - n} r_k^n. \tag{18}$$

The approximation is appropriate for sufficiently small values of $\rho$. Using this approximate formula, we can calculate the minimum synteny block size that will appear under gene order randomization with probability $1 - p_k(0) < \alpha$

$$k_{\min} = \left\lceil \frac{\log\left(1 - (1-\alpha)^{1/(2\tilde{N}_a\tilde{N}_b)}\right) - 2\log(1-\rho)}{\log \rho} \right\rceil \tag{19}$$

so that

$$k \geq k_{\min} \iff 1 - p_k(0) < \alpha \tag{20}$$

assuming the approximation in Eq. 18 holds. In practice, we round the right hand side of Eq. 19 to the nearest integer rather than using the ceiling operation. Alternatively, we can calculate the expected random fraction of nanosynteny blocks with $k$ genes, defined as the expected number of nanosynteny blocks with $k$ genes found under random permutations divided by the observed number of nanosynteny blocks with $k$ genes in the non-permuted homology matrix

$$\mathbb{E}[f_k] = \frac{2\tilde{N}_a\tilde{N}_b r_k}{\hat{n}_k}. \tag{21}$$

and pick $k_{\min}$ to be the largest $k$ satisfying

$$\mathbb{E}[f_k] < \alpha. \tag{22}$$

We further require that at least $k_{\min}$ distinct orthogroups are represented in a nanosynteny block.

#### 4.4 Step 3: Identify candidate pairs of nanosynteny blocks

Next, we created a directed acyclic graph of nanosynteny blocks for each chromosome pair in each pairwise species comparison. There is an edge from nanosynteny block $\mathbf{s}$ to $\mathbf{s}'$ if they are on the same pair of chromosomes, have the same slope, if

$$d_{\text{nano}}(\mathbf{s}, \mathbf{s}') \leq d_{\text{nano}}^{\max} \tag{23}$$

is satisfied, and if

$$x_A < x'_A \text{ and } x_B < x'_B \text{ for } \mathbf{x} \in \mathbf{s}, \mathbf{x}' \in \mathbf{s}' \tag{24}$$

for $\text{slope}(\mathbf{s}) = \text{slope}(\mathbf{s}') = 1$ or

$$x_A < x'_A \text{ and } x_B > x'_B \text{ for } \mathbf{x} \in \mathbf{s}, \mathbf{x}' \in \mathbf{s}' \tag{25}$$

for $\text{slope}(\mathbf{s}) = \text{slope}(\mathbf{s}') = -1$.

#### 4.5 Step 4: Decide connections between nanosynteny blocks

We then examine each pair of connected nanosynteny blocks to decide whether they should form a preliminary microsynteny block. We consider dots, $\mathbf{y} = (y_A, y_B)$, for potentially bridging two nanosynteny blocks if they are not in nanosynteny blocks and satisfy

$$x_A < y_A < x'_A \text{ and } x_B < y_B < x'_B \text{ for } \mathbf{x} \in \mathbf{s}, \mathbf{x}' \in \mathbf{s}' \tag{26}$$

for $\text{slope}(\mathbf{s}) = \text{slope}(\mathbf{s}') = 1$ or

$$x_A < y_A < x'_A \text{ and } x_B > y_B > x'_B \text{ for } \mathbf{x} \in \mathbf{s}, \mathbf{x}' \in \mathbf{s}' \tag{27}$$

for $\text{slope}(\mathbf{s}) = \text{slope}(\mathbf{s}') = -1$. We define the terminal dots of two neighboring nanosynteny blocks to be

$$\mathbf{y}_0 \in \mathbf{s}, \mathbf{y}_{m+1} \in \mathbf{s}' \text{ such that } d_H(\mathbf{y}_0, \mathbf{y}_{m+1}) = d_{\text{nano}}(\mathbf{s}, \mathbf{s}'). \tag{28}$$

We then search for eligible dots $\mathbf{y}_1, \mathbf{y}_2, \ldots, \mathbf{y}_m$ from which we can create a sequence $\mathbf{y}_0, \mathbf{y}_1, \mathbf{y}_2, \ldots, \mathbf{y}_{m+1}$ that is either monotonically increasing or decreasing in species $B$ indices as species $A$ indices are increased and for which the distance between any two sequential dots satisfies

$$d_H(\mathbf{y}_i, \mathbf{y}_{i+1}) \leq d_{\text{dot}}^{\max}. \tag{29}$$

If there is at least one such path between the nanosynteny blocks, we search for the longest such path, and merge the two nanosynteny blocks with the dots in the path into a microsynteny block. We repeat this with all candidate nanosynteny block pairs, noting that any number of nanosynteny blocks can potentially be combined into a single microsynteny block.

### 5 Permutation tests for nanosynteny

In order to test for the possibility of finding nanosynteny in homology matrices by random chance, we randomly permuted the rows and columns of each homology matrix and looked for nanosynteny within the permuted homology matrices. This process preserves the sizes of each orthogroup while scrambling the order of genes on each genome. We then calculated the distribution of nanosynteny block sizes (measured in number of TDEC pairs) for each permuted homology matrix, relative to the number of nanosynteny blocks of each size in the original homology matrix. We call this the random fraction of nanosynteny blocks, $f_k$, for blocks containing $k$ pairs of TDECs. We also calculated the expected random fraction $\mathbb{E}[f_k]$ (Eq. 21). The observed and expected random fractions of nanosynteny blocks were in close correspondence (Fig. S2), suggesting that one can find the minimum nanosyteny size $k_{\min}$ with respect to a significance cutoff $\alpha$ (Eq. 22) using only the expected random fraction calculated from the dimensions and density of the homology matrix along with the nanosynteny distribution of the original homology matrix.

#### 6 Inferring multi-gene duplications and their timings

See the Methods section in the main text.

#### 7 Analysis of gene distances

We used the normalized Levenstein distance as measure of sequence similarity between amino acid sequences, which measures the edit distance between two strings. This explicitly ignores all differences in amino acid substitution probabilities for a simple measure of substitutions per site.

The Levenshtein distances between two TDECs, $\tilde{g}^A_{ci}$ and $\tilde{g}^B_{c'j}$, is defined as the average over all pairs of genes across the tandem duplication blocks

$$d_L(\tilde{g}^A_{ci}, \tilde{g}^B_{c'j}) = \frac{1}{|\tilde{g}^A_{ci}||\tilde{g}^B_{c'j}|} \sum_{g \in \tilde{g}^A_{ci}} \sum_{g' \in \tilde{g}^B_{c'j}} d_L(g, g'). \quad (30)$$

Note that this formula reduces to the normal Levenshtein distance if the two TDECs are trivial (each contain a single gene).

#### 8 Inferring amino acid substitution rates

We use the generic tree from Fig. 4d, as it applies to all after-speciation duplications. We can determine amino acid substitution rates by calculating

$$\Delta d_L = d_L(A1, B1) - d_L(A2, B1) \quad (31)$$
$$= (r_{\text{source}} - r_{\text{target}})t_d \quad (32)$$

and

$$d_L(A1, A2) = (r_{\text{source}} + r_{\text{target}})t_d \quad (33)$$

and simply calculating the rates (in units of substitutions per site per MYA) as

$$r_{\text{source}} = \frac{d(A1, A2) + \Delta d_L}{2t_d} \quad (34)$$
$$r_{\text{target}} = \frac{d(A1, A2) - \Delta d_L}{2t_d}. \quad (35)$$

The expected number of substitutions on each branch after the duplication are

$$\lambda_{\text{source}} = r_{\text{source}} t_d L \tag{36}$$

$$\lambda_{\text{target}} = r_{\text{target}} t_d L \tag{37}$$

where $L$ is the number of amino acids in the protein (calculated as the average of the number of amino acids in A1 and A2). Using an infinite sites approximation, we can model the probability of finding $k$ substitutions in a branch according to the Poisson distribution

$$p(k;\lambda) = \frac{\lambda^k e^{-\lambda}}{k!}. \tag{38}$$

In order to rule out the possibility that our calculated rates are different due to random chance, we can calculate the probability of observing differences in the number of substitutions between A1-B1 and A2-B1 ($\frac{\Delta d_L L}{t_d}$)

$$q(\Delta k;\lambda_1,\lambda_2) = \sum_{k=\max(0,-\Delta k)}^{L+\min(0,-\Delta k)} p(k;\lambda_1)p(k+\Delta k;\lambda_2) \tag{39}$$

where $\lambda_1$ and $\lambda_2$ are the expected number of substitutions on the branches leading to A1, and A2 and $-L \leq \Delta k \leq L$.

The null hypothesis is that the observed $\Delta d_L$ is a result of evolution since the duplication event at either the source or target rate. Because the larger of the two rates will lead to a larger spread in the null model distribution, we only need to test against the larger rate, $\lambda_{\max} = \max(\lambda_{\text{source}}, \lambda_{\text{target}})$. We can calculate a $p$ value from the two-sided test

$$\begin{aligned} p &= q\left(\Delta K \geq \left|\frac{\Delta d_L L}{t_d}\right|; \lambda_{\max}, \lambda_{\max}\right) \\ &= \sum_{\Delta k=\left\lceil \frac{\Delta d_L L}{t_d}\right\rceil}^{L} q(\Delta k;\lambda_{\max},\lambda_{\max}) + \sum_{\Delta k=-\left\lceil \frac{\Delta d_L L}{t_d}\right\rceil}^{-L} q(\Delta k;\lambda_{\max},\lambda_{\max}) \end{aligned} \tag{40}$$

where $\frac{\Delta d_L L}{t_d}$ is the difference in the number of substitutions between the source-reference pair and the target-reference pair. We consider a difference in rates to be significant when $p \leq 0.05$.

### 9 Analysis of epigenetic data

We downloaded human ChIP-seq data from the ENCODE portal with the following identifiers: ENCFF003GBT, ENCFF003NRZ, ENCFF003PFV, ENCFF004JWB, ENCFF009ARK, ENCFF010BPD, ENCFF011PCV, ENCFF013JSB, ENCFF014HCV, ENCFF015KVF, ENCFF015TWY, ENCFF016YLR, ENCFF019GQD, ENCFF020IPM, ENCFF020XSD, ENCFF020XZP, ENCFF022BPF, ENCFF028VXO, ENCFF031POW, ENCFF031TEU, ENCFF032FEA, ENCFF032HKU, ENCFF033VFV, ENCFF034PBB, ENCFF037OSK,

ENCFF038UXA, ENCFF039NIU, ENCFF042SPE, ENCFF042STK, ENCFF044KLK, ENCFF047SBJ, ENCFF048ZUF, ENCFF050PLB, ENCFF051FKP, ENCFF051GOU, ENCFF051QIL, ENCFF054UNU, ENCFF054VRQ, ENCFF056BKL, ENCFF058QBP, ENCFF063INA, ENCFF069DWA, ENCFF081IBY, ENCFF083TRJ, ENCFF084TKR, ENCFF088IXP, ENCFF089KQW, ENCFF092VQF, ENCFF093WAF, ENCFF095ITH, ENCFF095XUU, ENCFF098UDL, ENCFF101XFJ, ENCFF105GJO, ENCFF106ZZM, ENCFF107XHS, ENCFF108LTK, ENCFF114HWE, ENCFF115EHH, ENCFF116TJK, ENCFF120RBR, ENCFF122LKO, ENCFF122QLT, ENCFF123SJE, ENCFF124NOK, ENCFF126HOZ, ENCFF126LKO, ENCFF128BBO, ENCFF128BKW, ENCFF130HHC, ENCFF130NUG, ENCFF132YWJ, ENCFF135UNI, ENCFF140GAT, ENCFF141USZ, ENCFF148JSK, ENCFF148LUF, ENCFF150SAI, ENCFF153OFF, ENCFF154LAJ, ENCFF157ZVJ, ENCFF163DHS, ENCFF163PTO, ENCFF169MHR, ENCFF173NSX, ENCFF174HPH, ENCFF174WIT, ENCFF174XHL, ENCFF179EUX, ENCFF180UYC, ENCFF181HLF, ENCFF182YXH, ENCFF183ZTW, ENCFF186PZN, ENCFF191CJN, ENCFF195NGS, ENCFF199GYV, ENCFF211RIL, ENCFF212KEL, ENCFF212ZFW, ENCFF215IDC, ENCFF225AOQ, ENCFF231VIR, ENCFF234TIE, ENCFF237BCW, ENCFF237SLF, ENCFF241MGV, ENCFF242OBC, ENCFF243OOK, ENCFF243TWF, ENCFF243ZOW, ENCFF246QNM, ENCFF249ILQ, ENCFF251CXB, ENCFF254FKJ, ENCFF257HZY, ENCFF257QES, ENCFF257SDT, ENCFF258EWK, ENCFF258MPY, ENCFF259GNO, ENCFF261VFS, ENCFF264ASR, ENCFF264HRZ, ENCFF264LWT, ENCFF264MFY, ENCFF266AHD, ENCFF266EAY, ENCFF266SCK, ENCFF267RUZ, ENCFF271URK, ENCFF271YEZ, ENCFF273JQE, ENCFF273KWS, ENCFF274UTN, ENCFF275OLK, ENCFF276UFE, ENCFF277VLD, ENCFF278QLF, ENCFF279GIO, ENCFF279SBT, ENCFF280QYJ, ENCFF282SRM, ENCFF282VQS, ENCFF284NUP, ENCFF287BWZ, ENCFF288QMN, ENCFF290HDV, ENCFF290QEI, ENCFF291UTT, ENCFF293EEX, ENCFF293OTU, ENCFF295YHH, ENCFF297LKI, ENCFF297SWX, ENCFF300XQV, ENCFF300YMQ, ENCFF303PLE, ENCFF309SJG, ENCFF311AYH, ENCFF313XUD, ENCFF319GFK, ENCFF320PWH, ENCFF321CUR, ENCFF321LZL, ENCFF321VNT, ENCFF323HEP, ENCFF325EMH, ENCFF331OKQ, ENCFF331SDA, ENCFF335QYK, ENCFF337EUB, ENCFF337EZM, ENCFF338HOP, ENCFF344RCN, ENCFF346JRR, ENCFF352AAW, ENCFF353GUK, ENCFF354FYC, ENCFF354VHC, ENCFF358QPI, ENCFF358RAF, ENCFF359DGK, ENCFF359FNY, ENCFF364CCZ, ENCFF370MEL, ENCFF371BOR, ENCFF371MIJ, ENCFF372NAI, ENCFF375YPQ, ENCFF376RDO, ENCFF378TNA, ENCFF380OMR, ENCFF386SPQ, ENCFF386WIJ, ENCFF388HBJ, ENCFF390SFV, ENCFF392RBS, ENCFF392VXP, ENCFF393ZPK, ENCFF397QED, ENCFF400KZY, ENCFF400NQI, ENCFF402CVL, ENCFF404DAA, ENCFF405NQM, ENCFF412JKW, ENCFF414FEQ, ENCFF414RVW, ENCFF417ZVA, ENCFF420MJH, ENCFF420RNX, ENCFF423YBA, ENCFF425FMW, ENCFF428HGT, ENCFF432TOV, ENCFF437CFO, ENCFF438TGS, ENCFF438UYU, ENCFF439PPA, ENCFF441IAJ, ENCFF441OEQ, ENCFF441XOS, ENCFF451OTN, ENCFF453NOK, ENCFF468UEP, ENCFF471BAS, ENCFF474STO, ENCFF477FDT, ENCFF478XEW, ENCFF480XEY, ENCFF485BWC, ENCFF485YEL, ENCFF489NKI, ENCFF491TUW,

ENCFF495CMD, ENCFF499HWN, ENCFF499QCN, ENCFF499XRO, ENCFF504DKP, ENCFF504XAW, ENCFF508OUM, ENCFF509VVM, ENCFF513OZY, ENCFF516GKI, ENCFF521BGK, ENCFF522EVU, ENCFF525LZQ, ENCFF528HJI, ENCFF529DXH, ENCFF529IHV, ENCFF534HLN, ENCFF538GPP, ENCFF539LWO, ENCFF541CWH, ENCFF542VXZ, ENCFF544BVR, ENCFF545KUT, ENCFF549EPL, ENCFF551GZK, ENCFF553OXO, ENCFF556ZLH, ENCFF557HHH, ENCFF559NXP, ENCFF562LJU, ENCFF563LPU, ENCFF566AVE, ENCFF568IBR, ENCFF569FQL, ENCFF569OIE, ENCFF571XOZ, ENCFF575QGX, ENCFF579DHC, ENCFF580KZB, ENCFF580ZNI, ENCFF581DIL, ENCFF583VWE, ENCFF584CKL, ENCFF586EPU, ENCFF586KGP, ENCFF587PUE, ENCFF599JQM, ENCFF608YXU, ENCFF612UQT, ENCFF615ZEK, ENCFF618WXG, ENCFF620NLZ, ENCFF621KKH, ENCFF623BIK, ENCFF623WKE, ENCFF624XUA, ENCFF628SPP, ENCFF631LYE, ENCFF634LYO, ENCFF636EZQ, ENCFF636MPX, ENCFF638ZNV, ENCFF639IIQ, ENCFF640NVF, ENCFF640WKU, ENCFF641PZC, ENCFF642MOF, ENCFF642TNR, ENCFF645DEG, ENCFF647MKA, ENCFF648NVI, ENCFF650EZM, ENCFF651EWX, ENCFF654BGS, ENCFF656CLB, ENCFF657OCV, ENCFF661CVI, ENCFF665CXY, ENCFF665TYI, ENCFF673DXP, ENCFF676ZJY, ENCFF680LKL, ENCFF683LBN, ENCFF685DRX, ENCFF688BBH, ENCFF690JQD, ENCFF692LNN, ENCFF693OWK, ENCFF694OEQ, ENCFF696UEY, ENCFF702ZYG, ENCFF703AQK, ENCFF703LZT, ENCFF703WUP, ENCFF707BCP, ENCFF707OIS, ENCFF709UON, ENCFF714QJF, ENCFF720AIQ, ENCFF722IPW, ENCFF725XRS, ENCFF726GVA, ENCFF729RVV, ENCFF730IAW, ENCFF733IXH, ENCFF733OPS, ENCFF733YQV, ENCFF734JTB, ENCFF735RJD, ENCFF738BQB, ENCFF740RYA, ENCFF740UPD, ENCFF744ADU, ENCFF762XSC, ENCFF768JKP, ENCFF772PDE, ENCFF773TCN, ENCFF774UOV, ENCFF779GTF, ENCFF780EJQ, ENCFF780EZC, ENCFF780OBE, ENCFF781ELK, ENCFF781JAZ, ENCFF782ZWK, ENCFF789HLD, ENCFF793HOY, ENCFF793YZF, ENCFF794WNF, ENCFF795QHB, ENCFF796MRD, ENCFF800QBA, ENCFF803DMD, ENCFF805RHU, ENCFF807WWP, ENCFF816LQH, ENCFF816ZXG, ENCFF817RMG, ENCFF825GHS, ENCFF828VTG, ENCFF829COF, ENCFF830YPS, ENCFF837JRN, ENCFF838MXD, ENCFF839CNL, ENCFF839JMI, ENCFF839NZT, ENCFF848SZN, ENCFF849YNY, ENCFF851AEJ, ENCFF852FGF, ENCFF852MOU, ENCFF857WKM, ENCFF859IVY, ENCFF860MMV, ENCFF863EJN, ENCFF863OBQ, ENCFF865CYY, ENCFF865ZUH, ENCFF868QXK, ENCFF870HKK, ENCFF871QOG, ENCFF872KPX, ENCFF872VOX, ENCFF873KYO, ENCFF875WOF, ENCFF877ROD, ENCFF879XVO, ENCFF881UNX, ENCFF884KHZ, ENCFF887LSD, ENCFF888YYK, ENCFF889NGP, ENCFF893TFN, ENCFF896KEC, ENCFF900HNB, ENCFF900SUK, ENCFF902HIA, ENCFF904YBG, ENCFF907XXC, ENCFF909EZR, ENCFF917QCC, ENCFF918JSW, ENCFF918SRL, ENCFF922PBC, ENCFF922YMQ, ENCFF928LLI, ENCFF934TJQ, ENCFF934WEQ, ENCFF935CUB, ENCFF938WJH, ENCFF940BVW, ENCFF940UMR, ENCFF943LXJ, ENCFF949GYG, ENCFF949TTW, ENCFF950LIB, ENCFF950NLS, ENCFF952ONE, ENCFF953BNG, ENCFF955DUI, ENCFF955MQX, ENCFF955WLM, ENCFF958EIZ, ENCFF960KLD, ENCFF962HWM, ENCFF963LIO, ENCFF967HEU, ENCFF969SUY, ENCFF970KBT,

ENCFF972YUU, ENCFF972ZHA, ENCFF973UOE, ENCFF976KIV, ENCFF978EHV, ENCFF980FUE, ENCFF981AHU, ENCFF985SAO, ENCFF986TXQ, ENCFF995LGE, ENCFF996ZNX, ENCFF999DTR, ENCFF999DTR.

We downloaded mouse ChIP-seq data from the ENCODE portal with the following identifiers: ENCFF010WPI, ENCFF012YLY, ENCFF014FXK, ENCFF053TIX, ENCFF059DUD, ENCFF059NRX, ENCFF061YPO, ENCFF064XDW, ENCFF087KHI, ENCFF096CRU, ENCFF110BJJ, ENCFF115MDG, ENCFF127UZI, ENCFF132HEC, ENCFF171NVN, ENCFF178ARM, ENCFF193SPQ, ENCFF204TMG, ENCFF218WLW, ENCFF223OGJ, ENCFF232DFR, ENCFF232EMK, ENCFF235MXD, ENCFF240JEC, ENCFF256NRL, ENCFF268ONA, ENCFF315URY, ENCFF321KJL, ENCFF366FOW, ENCFF369FIZ, ENCFF379MTD, ENCFF382KIS, ENCFF401UJV, ENCFF403UMI, ENCFF404DFP, ENCFF404HAV, ENCFF409OJE, ENCFF416KHY, ENCFF427ERT, ENCFF432GOU, ENCFF434ATJ, ENCFF460LBN, ENCFF473GLQ, ENCFF475QXA, ENCFF487DVO, ENCFF491JRL, ENCFF497QTK, ENCFF502NYV, ENCFF512IEQ, ENCFF516RZP, ENCFF519FAF, ENCFF521JOH, ENCFF522GHK, ENCFF523GNO, ENCFF526APR, ENCFF539MQF, ENCFF546GNK, ENCFF547SSI, ENCFF558IIX, ENCFF568ZKA, ENCFF574CPR, ENCFF575WWU, ENCFF584FIM, ENCFF586QBL, ENCFF587TLG, ENCFF590PWL, ENCFF598BQZ, ENCFF599BFJ, ENCFF603URB, ENCFF611MPQ, ENCFF626VBO, ENCFF639GUZ, ENCFF665RKA, ENCFF668KMK, ENCFF669GBT, ENCFF669SQZ, ENCFF672HAT, ENCFF686YOP, ENCFF689BVK, ENCFF691NZR, ENCFF698AUE, ENCFF699ERH, ENCFF700ZRI, ENCFF707OAY, ENCFF708XKV, ENCFF709LXJ, ENCFF726SHR, ENCFF741UWZ, ENCFF747DMA, ENCFF752ZUO, ENCFF754TWR, ENCFF767EJO, ENCFF775QFB, ENCFF780AOQ, ENCFF788SLF, ENCFF804DUE, ENCFF808TTR, ENCFF810OMG, ENCFF812GVN, ENCFF812ISP, ENCFF813FIB, ENCFF827CCJ, ENCFF833XFE, ENCFF862RRW, ENCFF868AVS, ENCFF872IVY, ENCFF876QGY, ENCFF881WJZ, ENCFF890KSS, ENCFF896KSI, ENCFF910NVL, ENCFF928EJF, ENCFF928SBC, ENCFF938OUM, ENCFF938XOE, ENCFF948SLR, ENCFF950DSF, ENCFF953UZI, ENCFF955MFI, ENCFF958AGA, ENCFF979BTU, ENCFF982LBU, ENCFF985QXX, ENCFF988JDG, ENCFF991OXE, ENCFF994VNL, ENCFF994XUU, ENCFF998FXE, ENCFF998FXE.

#### 10 Analysis of gene expression data

We used RNA expression data for the following human tissues: placenta, spleen, adrenal gland, ovary, upper lobe of left lung, heart left ventricle, pancreas, right lobe of liver, stomach, thyroid gland, peyer's patch, esophagus muscularis mucosa, esophagus squamous epithelium, gastrocnemius medialis, gastroesophageal sphincter, lower leg skin, omental fat pad, right cardiac atrium, sigmoid colon, subcutaneous adipose tissue, suprapubic skin, tibial nerve, transverse colon, breast epithelium, heart right ventricle, uterus, aorta, body of pancreas, colonic mucosa, heart, left cardiac atrium, left colon,

| Chromatin mark | Association with expression | Reference |
|---|---|---|
| H3K4me1 | + | [14, 15, 16] |
| H3K4me2 | + | [14] |
| H3K4me3 | + | [14, 15, 16] |
| H3K36me3 | + | [15, 16] |
| H3K27ac | + | [14, 17] |
| H3K79me1 | + | [17] |
| H3K9me3 | - | [15, 16] |
| H3K27me3 | - | [14, 15, 16] |

Table S1: List of chromatin marks we used in this study.

left lung, liver, lower lobe of left lung, lung, mucosa of descending colon, posterior vena cava, prostate gland, psoas muscle, right atrium auricular region, testis, thoracic aorta, right ventricle myocardium inferior, right ventricle myocardium superior, ascending aorta, camera-type eye, cardiac septum, cerebellum, diencephalon, frontal cortex, kidney, left lobe of liver, left ventricle myocardium inferior, left ventricle myocardium superior, lower lobe of right lung, mesenteric fat pad, metanephros, mucosa of gallbladder, occipital lobe, parietal lobe, sciatic nerve, skeletal muscle tissue, skin of body, spinal cord, temporal lobe, tongue, umbilical cord, upper lobe of right lung, urinary bladder, vagina.

We used RNA expression data for the following mouse tissues: heart, adrenal gland, gastrocnemius, left cerebral cortex, layer of hippocampus, forebrain, hindbrain, liver, midbrain.

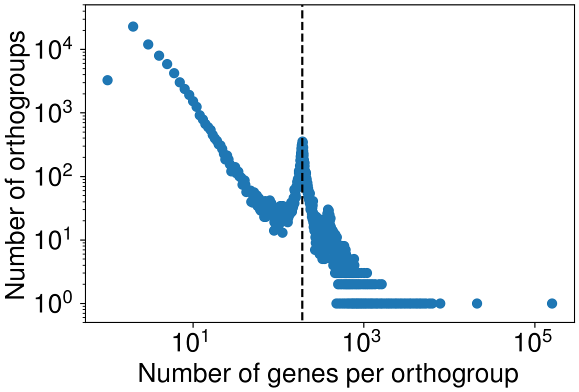


Figure S1: The distribution of orthogroup sizes, as inferred through OrthoFinder 2. Note dashed line corresponds to 194 genes in an orthogroup. The peak near 194 genes indicates that there are many orthogroups roughly corresponding to universal (in our dataset) single-copy orthologs. While there was a general trend of fewer orthogroups with large numbers of genes, there were stil a sizeable number of large (>> 194 genes) orthogroups which feature prominently in homology matrices.

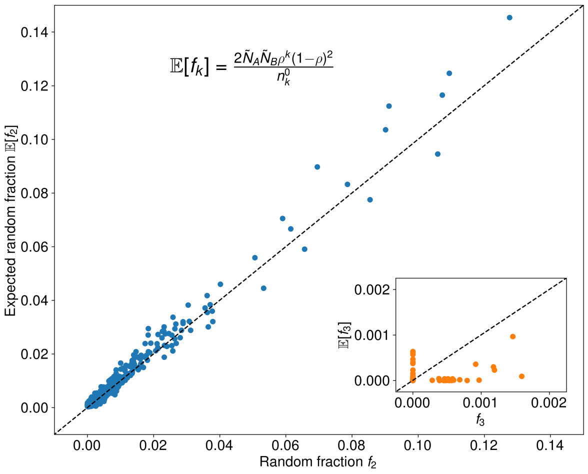


Figure S2: Correspondence between the observed random fraction of nanosynteny blocks and expected random fraction calculated through a percolation theoretic statistical model for all homology matrices. The main plot is for nanosynteny blocks containing two pairs of TDECs while the inset plot is for nanosyteny blocks with three pairs of TDECs. The close correspondence of observed and expected random fractions suggests that an estimate of the minimal nanosynteny block size $k_{\min}$ using the statistical model is justified.

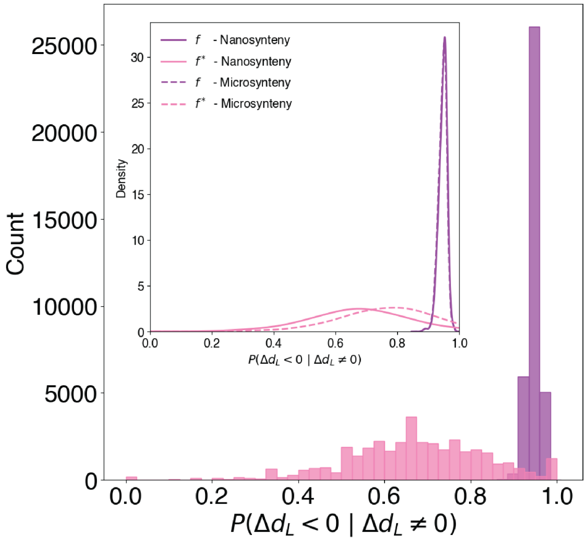


Figure S3: Nanosynteny preserves the reliability of microsynteny in predicting the most similar homolog in pairwise genome comparisons. The main plot presents the distributions of $f$ and $f^{\star}$, considering nanosynteny, for all 37,442 ordered pairs of species, excluding comparisons of genomes to themselves. In the inset plot we can see that the distribution of $f$ in the nanosynteny case (purple line) is sharply peaked around 0.9, as it's counterpart in microsynteny (pink line). On the other hand, the $f^{\star}$ distribution for the nanosynteny case (dotted purple line) has its peak slightly to the left than the microsynteny, without presenting significant differences.

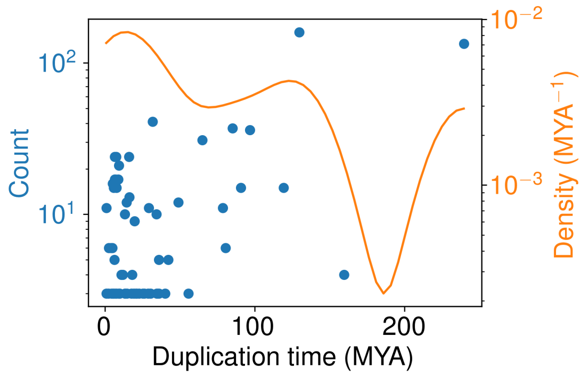


Figure S4: The distribution of inferred multi-gene duplication timings. We count each pair of duplicated regions in a species separately, ignoring the possibility of counting a single duplication event multiple times. The orange line is a Gaussian kernel density estimate, illustrating the high density of recent (smaller duplication time in MYA) duplications is comparable to older duplications.

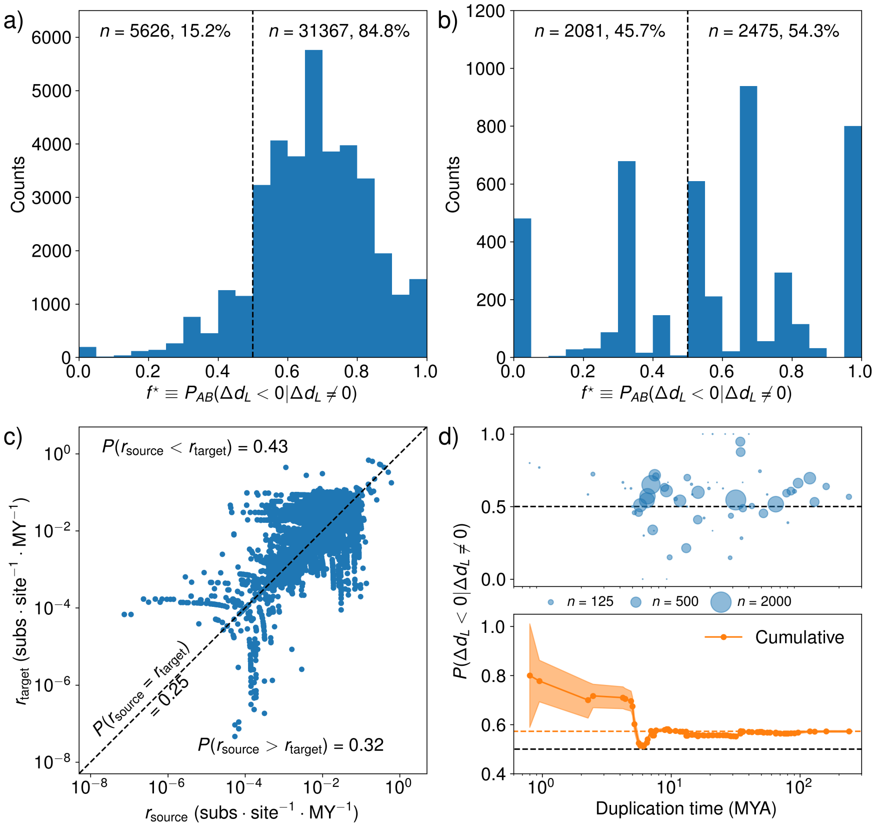


Figure S5: The results from Fig. 4 are insensitive to choice of microsynteny vs. nanosynteny. The results from Fig. 4 are reproduced here except using only nanosynteny, yielding very similar numbers for asymmetric evolution and polarization.

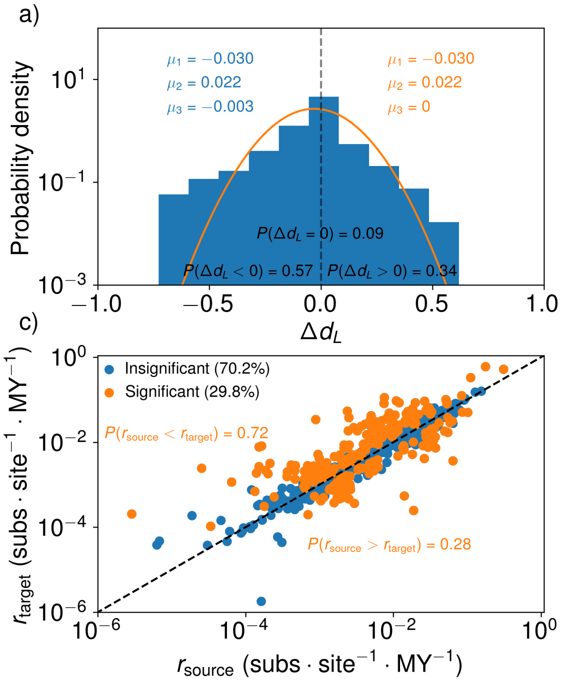


Figure S6: Asymmetric evolution and polarization of paralogs remains when averaging over reference orthologs, testing for significance of asymmetry, and restricting analysis to terminal duplications. a) The distribution of $\Delta d_L$ for all duplicated regions when averaging over reference orthologs. The orange line is a Gaussian distribution with the first two moments of the $\Delta d_L$ distribution. b) The distribution of $\Delta d_L$ for terminal duplicated regions when averaging over reference orthologs. c) Inferred substitution rates for all duplicated regions, averaged over reference orthologs. The orange dots represent cases where $r_{\text{source}}$ and $r_{\text{target}}$ are significantly different from one another. For significant cases, $P(r_{\text{source}} < r_{\text{target}}) = P(\Delta d_L \mid \Delta d_L \neq 0)$ is 0.72, as compared with the overall result of 0.593. d) The same analysis as for c), except with only terminal duplications.

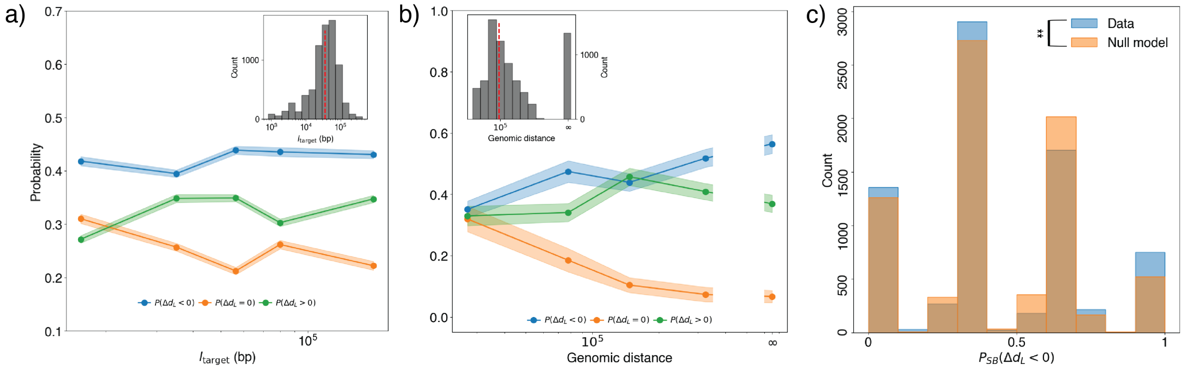


Figure S7: In nanosyteny, as in microsyteny (Fig. 5), the fate of large duplications is influenced by the size of the duplicated region and the genomic distance between copies and is shared for genes within a copy. a) The probability of the source copy evolving slower than the target copy, $P(\Delta d_L < 0)$, falls with larger target copy size in basepairs, $l_{\text{target}}$, while the probability of the source copy evolving faster, $P(\Delta d_L > 0)$, increases. The difference between the two probabilities represents the polarity of asymmetric paralog evolution, such that when $P(\Delta d_L < 0) < P(\Delta d_L > 0)$, the source copy is more likely to be the slowly evolving copy. Inset: the distribution of target copy sizes resulting from large duplications. b) When the same probabilities are examined as a function of the genomic distance between copies, both $P(\Delta d_L < 0)$ and $P(\Delta d_L > 0)$ increase with increased genomic distance. Again, the difference between the two probabilities indicates the polarity of asymmetric evolution. While there is not as clear of a trend as with $l_{\text{target}}$, at very small genomic distances $P(\Delta d_L < 0)$ and $P(\Delta d_L > 0)$ are very similar while at some larger distances $P(\Delta d_L < 0)$ and $P(\Delta d_L > 0)$ diverge. When the two copies are on different chromosomes (genomic distance of $\infty$), the difference between $P(\Delta d_L < 0)$ and $P(\Delta d_L > 0)$ is greatest. Note the monotonic decrease in $P(\Delta d_L = 0)$ with genomic distance. c) The probability of the source copy evolving slower than the target copy within a randomly chosen large duplication ($P_{SB}(\Delta d_L)$, $SB$ for synteny block) is significantly different from a null model produced by randomly resorting paralogous genes into new large duplications. If all paralogous gene pairs within a large duplicated region were polarized in evolutionary rate towards either the target or source copy, we would expect to see only values of $P_{SB}(\Delta d_L) = 0$ or $P_{SB}(\Delta d_L) = 1$. When compared with an ensemble of gene triplets randomly assigned to large duplications, the probability of observing our original data was low ($p < 0.05$), with excess counts at $P_{SB}(\Delta d_L) = 0$ and $P_{SB}(\Delta d_L) = 1$. This suggests that the fate of large duplications is determined collectively more often than expected by random chance.

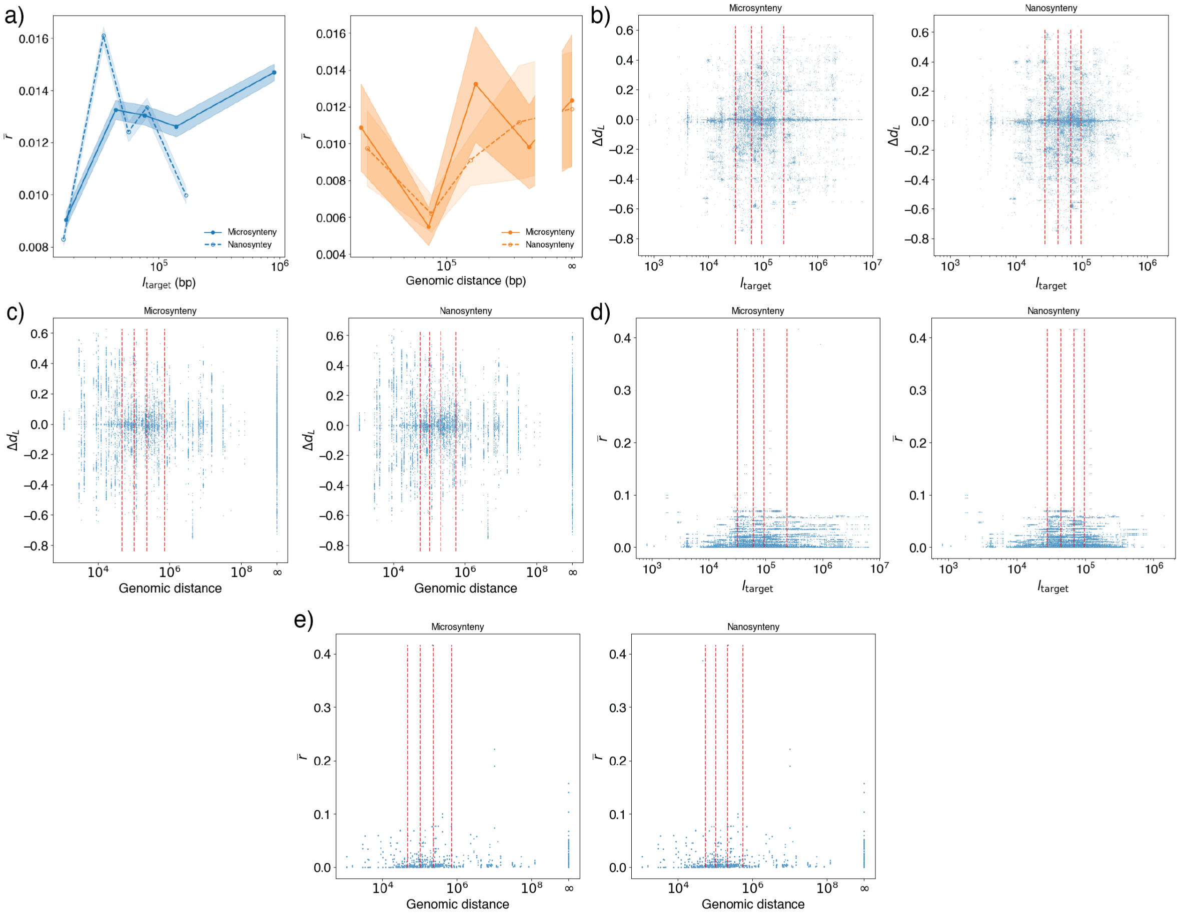


Figure S8: Nanosynteny vs. microsynteny comparison of the rough data to determine the fate of large duplications, size of duplicated region, genomic distance, and average amino acid substitution rate. a) Average amino acid substitution rate, $\overline{r}$, as a function of the size of the duplicated region, $l_{\mathrm{target}}$, (left) and genomic distance (right). It is easy to see that $\overline{r}$ does not have a clear trend that depends on any of the two size variables. b) Scatter plots of $\Delta d_L$ vs $l_{\mathrm{target}}$ and genomic distance for nanosyteny (right) and microsynteny (left). We use these scatter plots to compute the average behavior of $P(\Delta d_L > 0)$, $P(\Delta d_L < 0)$, $P(\Delta d_L = 0)$, each point in Fig. 5 a)-b) (microsynteny) and Fig. S7 a)-b) (nanosynteny), represents the average on each bin marked by the dotted red lines. c) Same as b) but for $\Delta d_L$ vs genomic distance. d) Same as b) but for $\overline{r}$ vs $l_{\mathrm{target}}$. e) Same as b) but for $\overline{r}$ vs genomic distance.

[9] DM Emms and Steven Kelly. STAG: species tree inference from all genes. *bioRxiv*, 2018.
